# A cell-ECM mechanism for connecting the ipsilateral eye to the brain

**DOI:** 10.1101/2021.03.11.434782

**Authors:** Jianmin Su, Yanping Liang, Ubadah Sabbagh, Lucie Olejníková, Ashley L. Russell, Jiang Chen, Yuchin Albert Pan, Jason W. Triplett, Michael A. Fox

## Abstract

Information about features in the visual world are parsed by circuits in the retina and are then transmitted to the brain by distinct subtypes of retinal ganglion cells (RGCs). Axons from RGC subtypes are stratified in retinorecipient brain nuclei, such as the superior colliculus (SC), to provide a segregated relay of parallel and feature-specific visual streams. Here, we sought to identify the molecular mechanisms that direct the stereotyped laminar targeting of these axons. We focused on ipsilateral-projecting subtypes of RGCs (ipsiRGCs) whose axons target a deep SC sublamina. We identified an extracellular glycoprotein, Nephronectin (NPNT), whose expression is restricted to this ipsiRGC-targeted sublamina. SC-derived NPNT and integrin receptors generated by ipsiRGCs are both required for the targeting of ipsiRGC axons to the deep sublamina of SC. Thus, a cell-extracellular matrix (ECM) recognition mechanism specifies precise laminar targeting of ipsiRGC axons and the assembly of eye-specific parallel visual pathways.

**Significance Statement:** Distinct features of the visual world are transmitted from the retina to the brain through anatomically segregated circuits. Despite this being an organizing principle of visual pathways in mammals, we lack an understanding of the signaling mechanisms guiding axons of different types of retinal neurons into segregated layers of brain regions. We explore this question by identifying how axons from the ipsilateral retina innervate a specific lamina of the superior colliculus. Our studies reveal a unique cell-extracellular matrix (ECM) recognition mechanism that specifies precise targeting of these axons to the superior colliculus. Loss of this mechanism not only resulted in the absence of this eye-specific visual circuit, but it led to an impairment of innate predatory visual behavior as well.

## Introduction

Parallel pathways encode, relay, and process information about distinct stimulus properties in all sensory systems. In the visual system, information about color, contrast, object motion, and light intensity are transmitted from the retina to brain in such parallel channels by retinal ganglion cells (RGCs). Over forty transcriptionally distinct subtypes of RGCs have been identified (1–5) and most project axons to different brain nuclei or even different regions within the same nuclei (5–10). In brain regions that process image-forming visual information, which in rodents includes the superior colliculus (SC) and dorsal lateral geniculate nucleus (dLGN), projections from distinct subtypes of RGCs are segregated into discrete sublamina. Despite this segregation being an organizing principle of parallel visual pathways in mammals, we lack an understanding of the molecular mechanisms underlying lamina-specific axon targeting.

To identify mechanisms that drive the laminar targeting of RGC axons, we focus here on the SC, the largest retinorecipient nucleus in rodents and a region responsible for driving goal-directed eye movements and a subset of innate visual behaviours (11, 12). In rodents, the majority of RGCs project axons to SC, where they arborize into the superficial-most domain in a subtype-specific fashion (5, 8). Transgenic tools labeling individual subtypes of RGCs have been instrumental in identifying lamina-specific projections from distinct RGC subtypes: for example, RGCs that convey information about object movement and direction selectivity project axons to the most superficial sublamina of SC, while ⍰RGCs project to deeper SC sublamina (9, 13–19). While the development of these transgenic tools only recently shed light on subtype-specific projection patterns, it has long been appreciated that axons from RGCs in the contralateral eye and ipsilateral eye (contraRGCs and ipsiRGCs, respectively) are targeted to distinct sublamina of SC (**Figure 1A&B**) (19, 20). This segregation of eye-specific inputs is important for coordinating coherent representations of the visual field from both eyes and is considered an essential building block of binocular vision (21).

**Figure 1.**
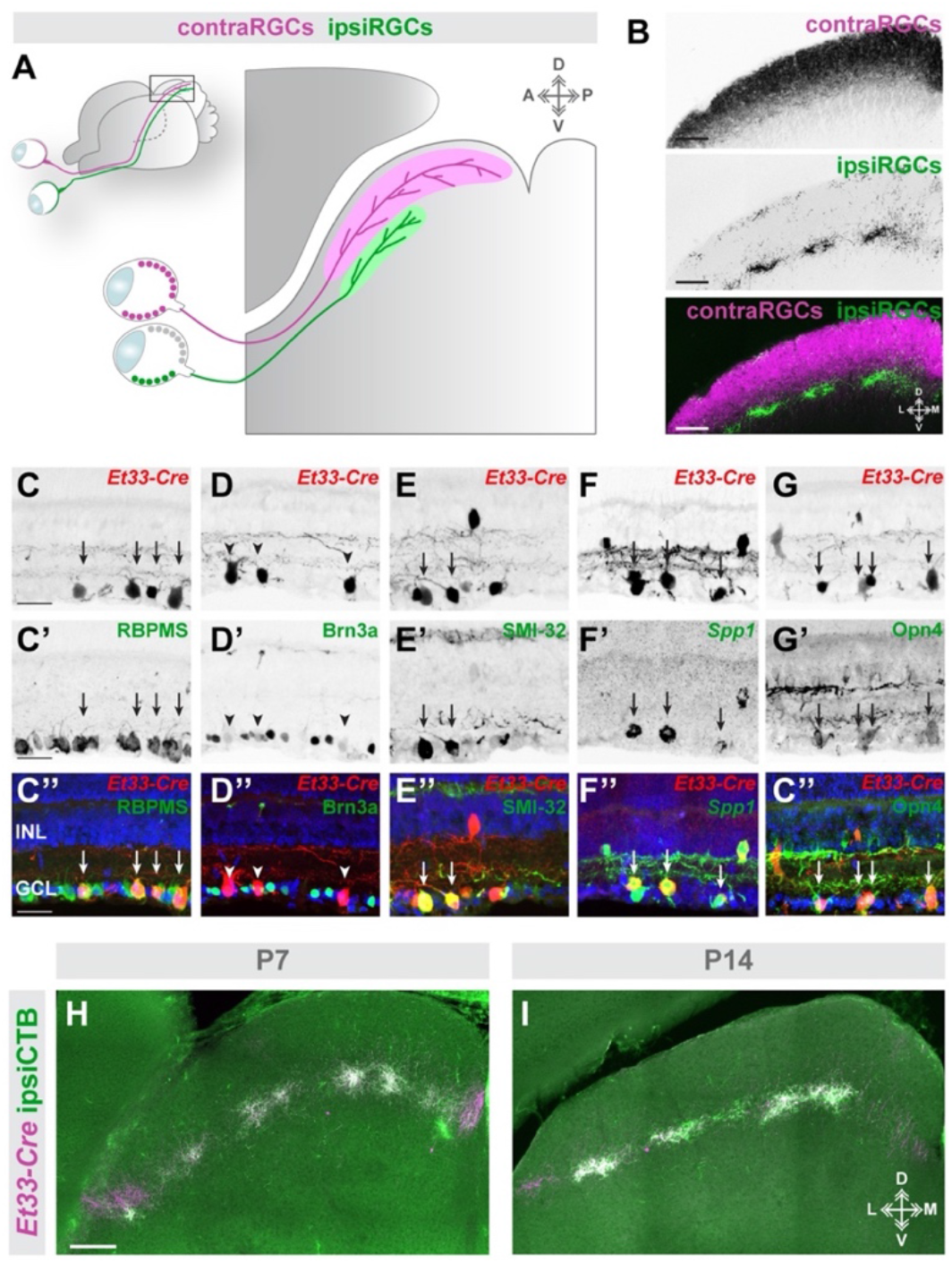
ipsiRGCs innervate the deepest sublamina of retinorecipient SC. **A.** Schematic representation of eye-specific retinocollicular projections. **B.** Coronal section depicting ipsi- and contraRGC projections to the P14 mouse SC labeled by intraocular delivery of different variants of fluorescently-conjugated CTB. **C-G.** Retinal cross sections from P12 *Et33-Cre::Rosa-Stop-tdT* mice in which RGCs are labeled by IHC (**C-E,G**) and ISH (**F**). RBMPS labels all RGCs; Brn3A labels contraRGCs; SMI32 and *Spp1* label subsets of αRGCs; Opn4 (Melanopsin) labels ipRGCs. Arrows highlight co-expression. Arrowheads depict tdT^+^ RGCs that do not express Brn3a. DAPI is shown in blue. **H,I**. Genetically labeled ipsiRGC axons target the deepest sublamina of the neo- and postnatal SC. Coronal sections of SC show tdT^+^ RGC axons in P7 and P14 *Et33-Cre::Rosa-Stop-tdT* mice contain fluorescently-conjugated CTB that was monocularly delivered into the ipsilateral eye. Scale bar in **B** = 250 μm; in **C** = 50 μm for **C-G**; in **H** = 250 μm for **H,I**.

While axons from some subtypes of RGCs initially overshoot their targets or transiently innervate inappropriate brain regions during development (22, 23), axons from ipsiRGCs initially target the appropriate sublamina of the SC (24). This suggests a selective developmental mechanism drives the laminar targeting of ipsiRGC axons and generates a segregated, eye-specific parallel pathway. Here, we identified a molecularly specified ECM ligand / cell surface receptor mechanisms that patterns ipsiRGC axon targeting to the SC. Spatially-restricted expression of the ECM protein Nephronectin (NPNT) is sufficient to promote the selective outgrowth of ipsiRGC axons *in vitro* and necessary for ipsiRGC axon targeting of SC *in vivo*. NPNT signals through RGD-dependent integrins and disrupting integrin signaling in ipsiRGCs (genetically or pharmacologically) impaired ipsiRGC axon growth on NPNT *in vitro* and impaired ipsiRGC innervation of SC *in vivo*. Taken together, these results shed light on a molecular matching mechanism that specifies laminar targeting of axons from the ipsilateral retina and establishes an eye-specific, parallel visual pathway.

## Results

### Ipsilateral projecting RGCs innervate a distinct sublamina of mouse superior colliculus

To label eye-specific RGC arbors in the developing SC, we delivered different fluorescently conjugated versions of cholera toxin subunit B (CTB) into each eye. Arbors of ipsiRGCs were confined to the anterior-most half of the SC and were in a deeper sublamina than arbors from contraRGCs (**Figure 1B**). Distinct projection patterns are not the only features that differentiate ipsiRGCs and contraRGCs; in fact, these subsets of RGCs are transcriptionally distinct (25) and can therefore be distinctly labeled transgenically. Here, we show that ipsiRGCs are specifically labeled in the *Et33-Cre::Rosa-Stop-tdT* mice (26). Not only is tdT expression restricted to the ventrotemporal crescent of retina, but all genetically labeled cells all co-express RBPMS (a marker of RGCs) and lack Brn3a (a marker for contraRGCs) (**Figure 1C&D**) (25, 27, 28). Note that tdT^+^cells present in the inner-most portion of the inner nuclear layer (INL) also express RBPMS, indicating they are displaced RGCs. This suggested that ipsiRGCs may themselves include several distinct subtypes. Indeed, labeling *Et33-Cre::Rosa-Stop-tdT* retina revealed that ipsiRGCs co-label with molecular markers associated with both ⍰RGCs (e.g. SMI-32 and *Spp1*) and intrinsically photosensitive RGCs (ipRGCs; e.g. melanopsin [Opn4])(**Figure 1E-G**). As expected, central projections of *Et33-Cre::Rosa-Stop-tdT*-labeled cells targeted brain regions known to be innervated by ipsiRGCs (as well as αRGCs and ipRGCs), including the suprachiasmatic nucleus (SCN), ventral lateral geniculate nucleus (vLGN), intergeniculate leaflet (IGL), dLGN, and SC (**Figure S1**). Anterograde labeling of ipsiRGC projections with CTB revealed that the majority, if not all, ipsiRGC projections to SC are labeled with tdT in neonatal and early postnatal *Et33-Cre::Rosa-Stop-tdT* mice. This analysis revealed a unique distinction in the patterns of retinorecipient innervation by ipsiRGCs. In the perinatal dLGN, ipsiRGC axons arborized broadly throughout the dLGN and were later refined to eye-specific domains by eye-opening (**Figure S1**)(29). In contrast, CTB^+^ /tdT^+^ ipsiRGC projections initially target the appropriate SC sublamina, rather than arborizing throughout the entire SC and then being refined into this sublamina (**Figure 1H&I**; see also (24)).

### Identification of a spatially-restricted ECM molecule in retinorecipient SC

Laminar targeting of axons could result from a specified molecular matching mechanism or from activity-dependent mechanism (which could include refinement or local growth). The initial specificity with which ipsiRGC axons target a segregated sublamina of SC suggested to us that a genetically specified matching mechanism might underlie the assembly of eye-specific visual pathways. What might this molecular matching mechanism be? To answer this question, we screened the Allen Brain Atlas (30) seeking to identify cell adhesion molecules, growth factors, morphogens, or extracellular matrix proteins whose expression were restricted to deep sublamina of the retinorecipient SC. One can didate that emerged was Nephronectin (NPNT), an ECM glycoprotein that contains epidermal growth factor-like repeats and multiple integrin binding motifs (31, 32). *In situ* hybridization revealed *Npnt*, the ge ne that encodes NPNT, was absent from most regions of the neonatal and postnatal mouse brain, with two notable exceptions: a deep sublamina of SC and two distinct layers of neocortex (**Figure 2A and S2**). Anterograde labeling of eye-specific RGC projections with CTB revealed that *Npnt^+^* cells were confined to the SC sublamina innervated by ipsiRGCs (**Figure 2B&C**). Importantly, NPNT protein was also generated by cells expressing *Npnt* transcripts and was enriched in a single lamina of the developing SC (**Figure 2D**).

**Figure 2.**
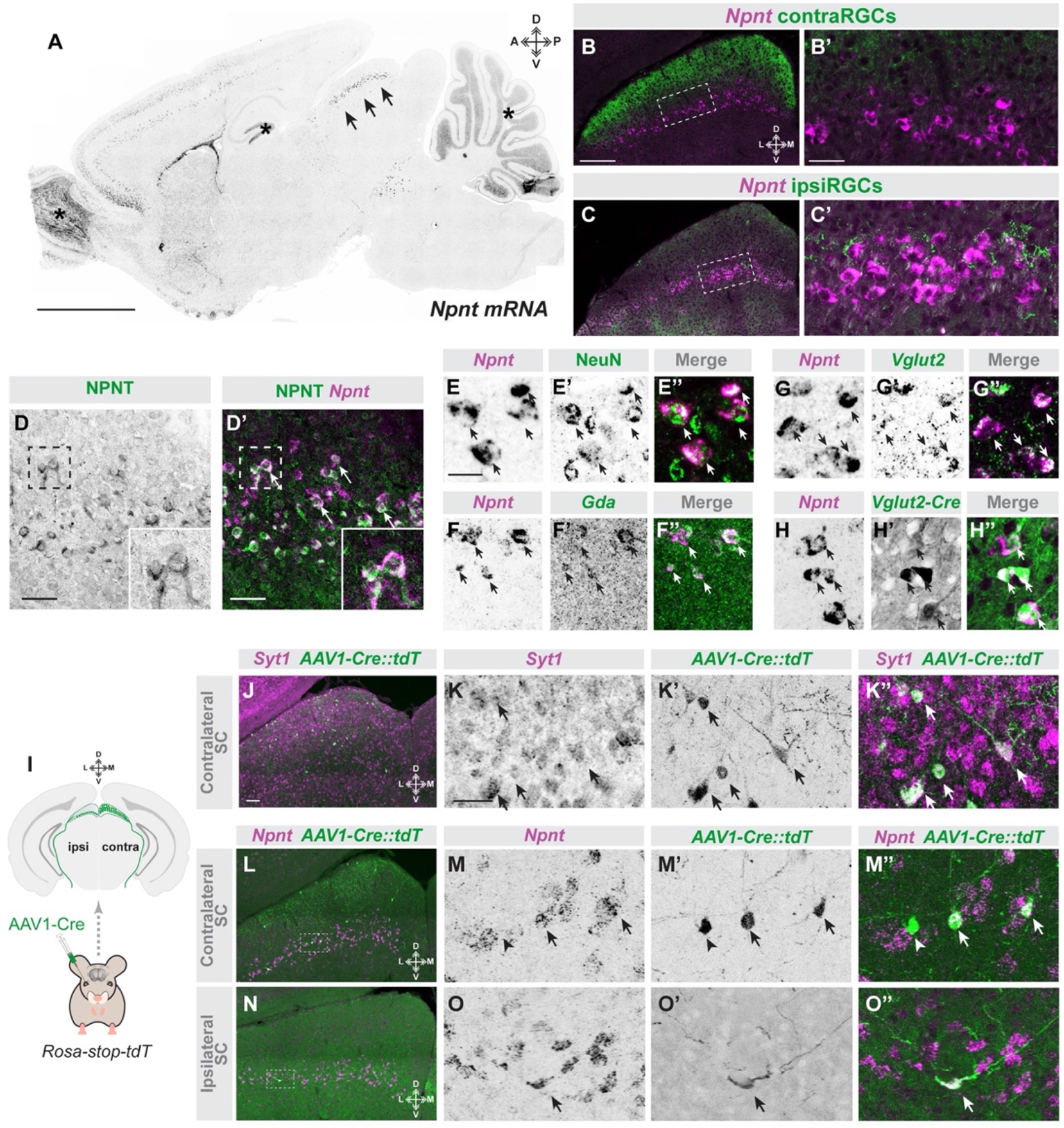
Nephronectin is generated by retino-recipient neurons in a restricted sublamina of the developing SC. **A.** ISH shows *Npnt* mRNA expression in a sagittal section of P8 mouse brain. Arrows highlight laminar expression of *Npnt* in SC. Asterisks denote brain region with high background signal which does not reflect true *Npnt* mRNA expression. **B,C.** ISH for *Npnt* mRNA in coronal section from P14 mouse brains in which contraRGC (**B**) and ipsiRGC (**C**) projections are labeled with CTB. **B’** and **C’** represent high magnification images of areas within dashed boxes in **B** and **C**, respectively. **D.** ISH for *Npnt* mRNA and IHC for NPNT protein in P14 SC. Insets in **D** and **D’** show high magnification images of *Npnt^+^* cells that generate NPNT protein. **E-H**. Excitatory neurons generate *Npnt*. ISH for *Npnt* mRNA was combined with IHC for NeuN (**E**) or ISH for *Gda* (**F**) or *Vglut2* (**G**) in P14 SC. **H** shows ISH for *Npnt* in the SC of *Vglut2-Cre::Thy1-stop-yfp* mice. **I**. Schematic depiction of monocular administration of AAV1-Cre to trans-synaptically label retino-recipient cells innervated by ipsi- or contraRGCs. **J,K.** SC cells labeled following monocular injection of AAV1-Cre in *Rosa-Stop-tdT* mice express neuronal *Syt1* mRNA. **K** depicts a high magnification image showing expression of *Syt1* mRNA in *AAvl-Cre::Rosa-Stop-tdT* (*AAV1-Cre::tdT*)-labeled cells.

We next sought to identify what cells generate *Npnt*. We probed *Npnt* mRNA expression in combination with molecular and genetic approaches that label neuronal and glial cell types in the developing SC (**Figure 2 & S2**). Expression of *Npnt* was restricted to a subset of excitatory neurons that co-expressed NeuN, *Syt1, Vglut2*, Calb, and *Gda1* (**Figure 2E-H & S2**). While NeuN, *Syt1, Vglut2*, and Calb were all found in larger subsets of neurons than those that generate *Npnt*, all *Gda1^+^* cells co-expressed *Npnt* and therefore also appeared to represent a small subset of SC neurons confined to the deepest sublamina of retinorecipient SC (**Figure 2**). *Npnt* expression was absent from every other cell type including astrocytes, microglia, *Gad1^+^*inhibitory neurons, *Sst* neurons, and *Pvalb^+^* neurons (**Figure S2**).

Knowing that *Npnt^+^* neurons are excitatory neurons (and most likely principal neurons in SC) we next asked whether these *Npnt^+^* neurons are themselves retinorecipient. To test this, we used a Cre-expressing adeno-associated virus (AAV1-Cre) which delivers Cre anterogradely across a single synapse (33–35). Intraocular delivery of AAV1-Cre into *Rosa-Stop-tdT* mice reliably labels *Syt1^+^* retinorecipient neurons in SC (**Figure 2I-K’’**). Using a monocular delivery approach, we determined that *Npnt\** neurons received direct input from ipsiRGCs (as well as from contraRGCs) (**Figure 2L-O’’**). Taken together, the structural properties of NPNT, the restriction of *Npnt^+^* neurons to the sublamina of SC innervated by ipsiRGC axons, and the synaptic connections between ipsiRGCs and *Npnt*^+^ cells, all suggest that NPNT may act as an ECM-based axonal targeting cue for ipsiRGCs.

### NPNT selectively promotes the growth of ipsiRGC axons

Despite having important roles in kidney development, whether or how NPNT influences neuronal and axonal development remains unexplored. Structurally related extracellular glycoproteins have well established roles in directing axonal guidance and targeting by their ability to bind and signal through receptors on the cell surface of the axonal growth cone. To test whether NPNT was similarly able to promote the growth of RGC axons, we turned to *in vitro* assays. RGCs represent an exceptionally small fraction (<1%) of all retinal cells, therefore, we purified RGCs from dissociated retinas by immunopanning and cultured these RGCs *in vitro* for several days (**Figure 3A & S3**). We hypothesized that NPNT may selectively promote the outgrowth of ipsiRGC axons based on its developmental expression pattern, therefore, we immunopanned RGCs from neonatal *Et33-Cre::Rosa-Stop-tdT* mice. This allowed us to generate diverse cultures that not only contained all subtypes of RGCs but contained genetically-labeled ipsiRGCs that were identifiable based on tdT expression (**Figure S3**).

**Figure 3.**
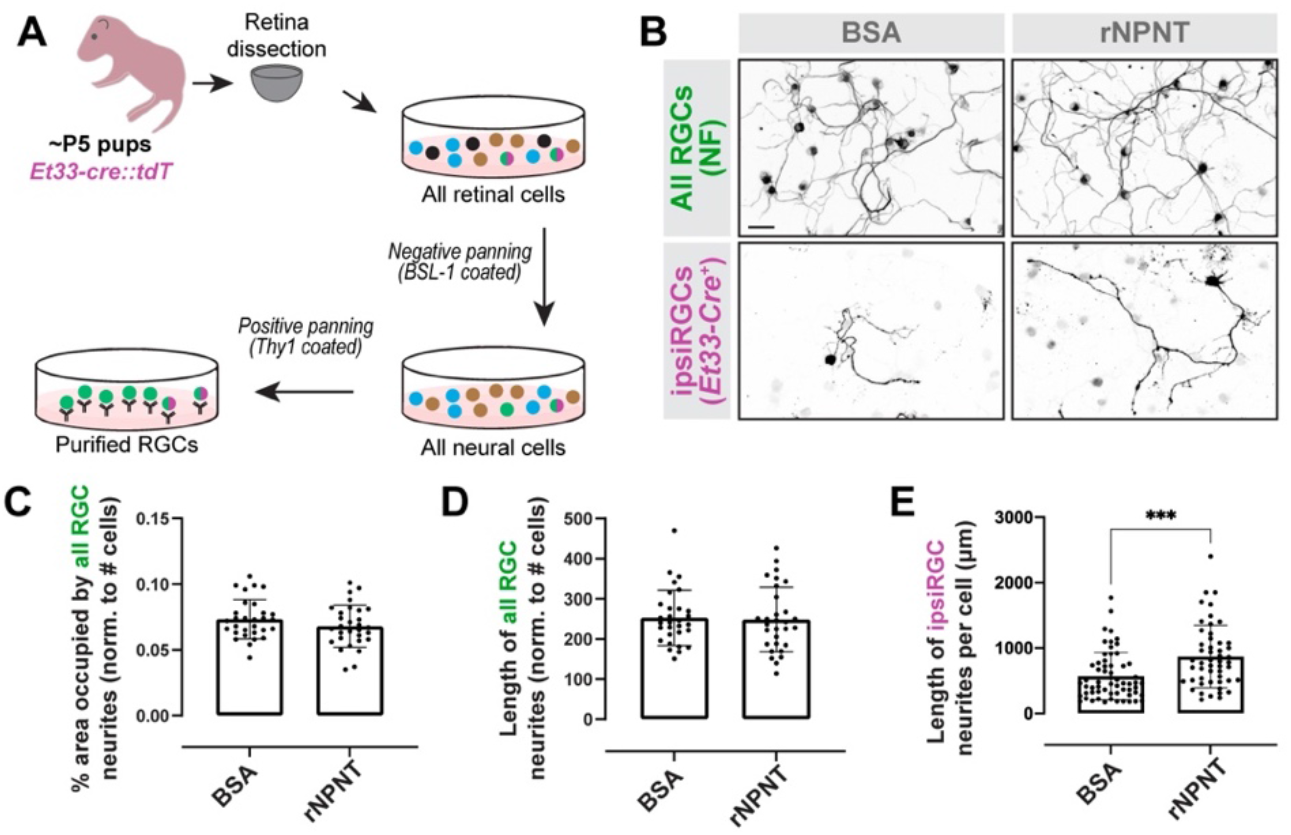
Nephronectin promotes ipsiRGC axon outgrowth via integrin signaling. **A.** Schematic representation of immunopanning RGCs from neonatal *Et33-Cre::Rosa-Stop-tdT* mice. **B**. RGCs from *Et33-Cre::Rosa-Stop-tdT* mice were cultures for 5 days on recombinant NPNT (rNPNT) or control substrates (BSA). All RGCs were labeled by IHC for neurofilament (NF). ipsiRGCs were identified by tdT expression (*Et33-Cre^+^*).**C,D**. Neurite outgrowth of all subtypes of RGCs from **B** was quantified by assessing the % area occupied by NF^+^ neurites or by measuring the total length of NF^+^ neurites per field of view. Data were normalized to the number of RGCs per field of view. Bars represent mean +/− SD. Data points represent a single field of view from a total of three experiments. ANOVA analysis indicated no significant differences between neurite outgrowth on these substrates. **E.** Neurite outgrowth of tdT^+^ ipsiRGCs from **B** was quantified by measuring the total length of tdT^+^ neurites. Bars represent mean +/− SD. Data points represent neurites from a single tdT^+^ ipsiRGC from a total of three experiments. P< 0.001 by ANOVA (n=61 cells in BSA group and 51 cells in rNPNT group). Scale bar in B = 50 μm and in F = 50 μm.

In order to test whether NPNT impacts the outgrowth of RGC axons, immunopanned RGCs were cultured on substratum containing recombinant NPNT (rNPNT) or control protein (BSA). After 5 days, cultures were fixed and immunostained with antibodies against neurofilament (NF) to label the neurites of all RGCs. The length of NF^+^ neurites was similar between RGCs grown on rNPNT and control substratum, suggesting NPNT had little impact on most RGC axons (**Figure 3B-D**). In contrast, when the length of tdT^+^ ipsiRGC neurites was measured, there was a significant increase in neurite length compared with control conditions (**Figure 3B&E**). This suggests that NPNT specifically promotes ipsiRGC axonal growth.

### NPNT is necessary for ipsiRGC axon targeting of SC

To test whether NPNT is required for retinocollicular circuit formation *in vivo*, we used a conditional allele to delete *Npnt (Npnt^fl/fl^*) from select neuronal populations (to avoid its necessity in kidney development; (36)). Based on the expression of *Npnt* by excitatory neurons in SC, we used two driver lines to delete *Npnt* expression from SC: *Nes-Cre* and *Vglut2-Cre* (**Figure S4**). The resulting mutants were viable and fertile and the loss of NPNT from *Nes^+^* and *Vglut2^+^* cells had little impact on gross brain morphology or the cytoarchitecture of the retina (**Figure S4**). Moreover, conditional loss of *Npnt* in *Npnt^fl/fl^::Nes-Cre* (*Nes-cKO*) and *Npnt^fl/fl^::Vglut2-Cre* (*Vglut2-cKO*) mutants did not alter the laminar distribution of *Gda^+^* neurons in SC, demonstrating that NPNT was dispensable for the appropriate distribution of cells in the sublamina of SC targeted by ipsiRGC axons (**Figure S4**).

To assess eye-specific axon targeting of SC, RGC projections were anterogradely labeled by monocular injections of CTB. The loss of NPNT in both *Npnt^fl/fl^::Nes-Cre* and *Npnt^fl/fl^::Vglut2-Cre* had little impact on the innervation of SC by contraRGCs (**Figure 4A&B**). In contrast, the loss of NPNT in these mutants resulted in dramatic loss of ipsiRGC axons in SC (**Figure 4A,C & S4**). It is noteworthy that ipsiRGC axon innervation of other retinorecipient regions, such as pretectal nucleus and visual thalamus (which do not generate *Npnt* in wildtype mice and which are innervated by the same cohorts of RGCs that innervate SC (5, 19)), is not altered in the absence of NPNT (**Figure 4A,D & E-G**). This suggests that the absence of ipsiRGC axons in the SC of these mutants is not due to a loss of ipsiRGCs. Moreover, not only do contraRGCs projections appear normal in NPNT mutants, so do the projections from individual subtypes of contraRGCs, such as the ON-OFF direction selective RGCs (oodsRGCs) labeled in *Trhr-GFP* mice (**Figure 4H&I**)(Rivlin-Etzion et al. 2011). We interpret these results to suggest that SC-derived NPNT is the extracellular matrix (ECM) recognition mechanism that specifies precise laminar targeting of ipsiRGC axons and the assembly of eye-specific visual pathways.

**Figure 4.**
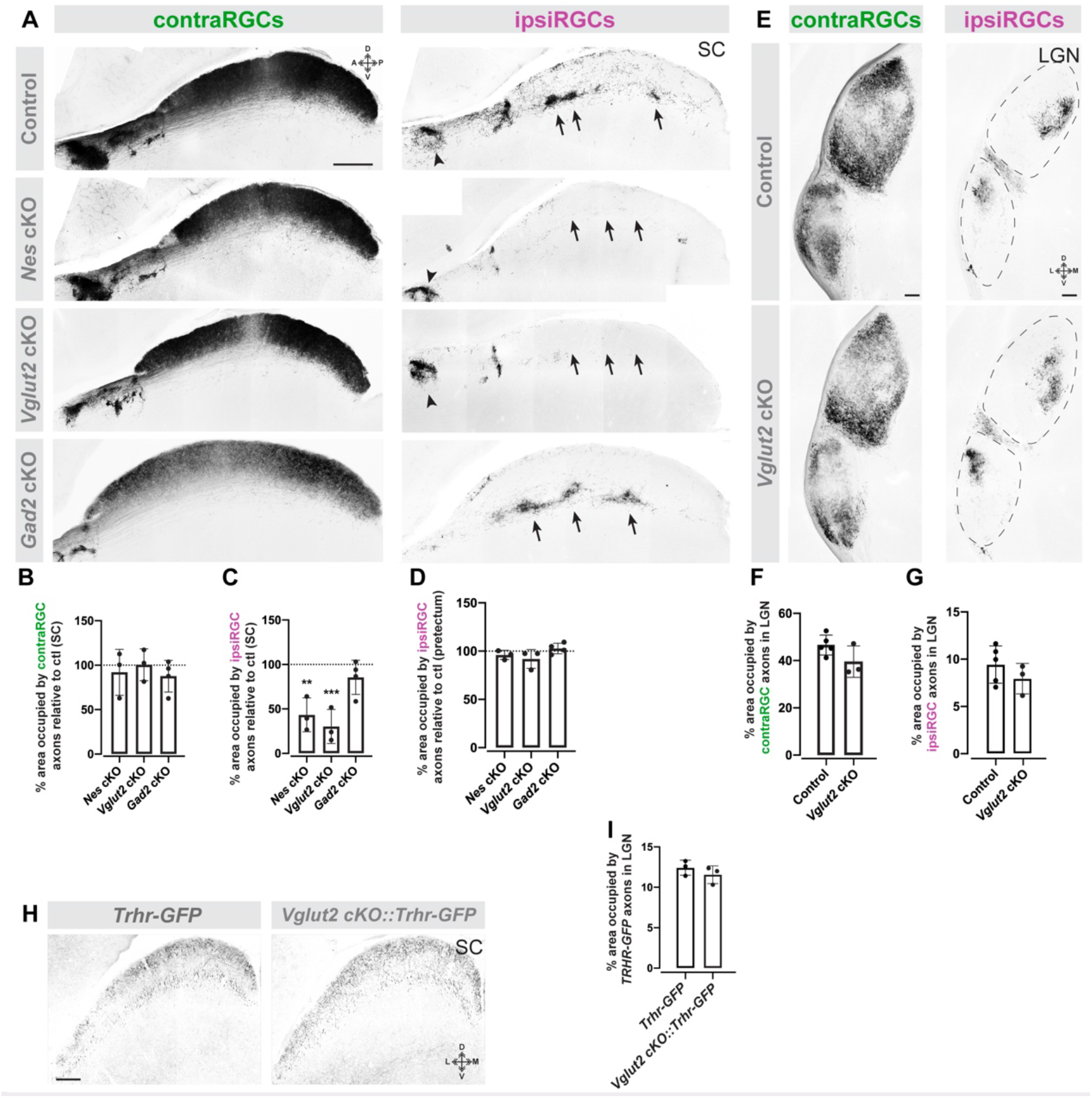
Nephronectin is required for ipsiRGC axon innervation of SC. **A.** CTB-labeled contra- and ipsiRGCs projections to SC in P14 control, *Npnt^fl/fl^::Nes-Cre* (*Nes* cKO); *Npnt^fl/fl^::Vglut2-Cre* (*Vglut2* cKO), and *Npnt^fl/fl^::Gad2-Cre* (*Gad2* cKO) mice. Images depict sagittal sections of SC. Arrows highlight sublamina of SC targeted by ipsiRGC axons in control and *Gad2* cKO mice, which are largely absent from *Nes* cKO and *Vglut2* cKO. Arrowheads highlight ipsiRGC axon targeting of pretectal nuclei. **B, C.** Quantification of the area of SC occupied by contra- (**B**) and ipsiRGC (**C**) projections in **A** compared to age-matched controls (represented by the dashed line). Bars represent means +/− SD. ** indicates P<0.005 and *** indicates P<0.0005 when compared to control by Student’s t-test (n=3 mice). **D.** Quantification of the area of pretectum occupied by ipsiRGC projections in **A** compared to age-matched controls (represented by the dashed line). Bars represent means +/− SD. **E.** CTB-labeled contra- and ipsiRGCs projections to visual thalamus (LGN) with CTB in P14 control and *Npnt^fl/fl^::Vglut2-Cre* (*Vglut2* cKO) mice. Dashed line encircle dLGN and vLGN. **F,G.** Quantification of the area of LGN occupied by contra- (**F**) and ipsiRGC (**G**) projections in **E** compared to age-matched controls (represented by the dashed line). Bars represent means +/− SD. **H.** On-Off direction selective RGC projections (in *Trhr-GFP* mice) appear unaltered in the absence of NPNT. Coronal sections from P14 *Trhr-GFP* control mice of *Npnt^fl/fl^::Vglut2-Cre::Trhr-GFP* (*Vglut2-cKO::Trhr-GFP*).**I.** Quantification of the % area of SC occupied by GFP^+^ projections in **H**. Bars represent means +/− SD. Scale bar in **A** = 500 μm, in **E** = 100 μm, and in **H** = 250 μm.

Since *Npnt* is expressed in subsets of amacrine cells and by infrequent cells in the ganglion cell layer (but not by ipsiRGCs labeled in *Et33-Cre::Rosa-Stop-tdT* retina or by many RGCs labeled in *Vglut2-Cre::Rosa-Stop-YFP* retina; **Figure S4**), it is also possible that retinal-derived NPNT contributes to the mistargeting of ipsiRGCs in *Npnt^fl/fl^::Nes-Cre* and *Npnt^fl/fl^::Vglut2-Cre* mutants. To rule out the role of retina-derived NPNT in ipsiRGC targeting of SC, *Npnt^fl/fl^* mice were crossed to *Gad2-Cre* (where Cre is expressed in amacrine cells, as well as *Npnt^-^* GABAergic neurons in SC), *Calb2-Cre* (which is expressed in >90% of RGCs; (37)), and *Et33-Cre*. In all three cases, the loss of NPNT from amacrine cells and/or RGCs had no significant impact on the targeting of ipsiRGCs to SC (**Figure 4A-C,7E & S4**). These results further demonstrate the specificity and selectivity that SC-derived NPNT plays in the precise laminar targeting of ipsiRGC axons.

### Integrin signaling is required for ipsiRGC targeting of SC

What might be the cell surface receptor on ipsiRGC axons that recognizes SC-derived NPNT and regulates the laminar targeting of these axons? The region of retina that contains ipsiRGCs (the ventrotemporal crescent) requires integrin binding for retinal cell adhesion to the ECM and neurite outgrowth *in vitro* (38). Since NPNT was initially identified in a screen for ligands of the RGD-binding α8β_1_ integrin in the embryonic kidney (31, 39), this suggests that a selective NPNT-integrin signaling mechanism may underlie axonal outgrowth and targeting of ipsiRGCs. To test this, we took a pharmacological approach to block integrin signaling and we re-assessed the ability of rNPNT to promote ipsiRGC axon outgrowth *in vitro*. Integrin signaling was blocking by treating immunopanned RGCs with integrin-blocking RGD peptides. Blocking RGD-dependent integrins in these assays reduced the ability of rNPNT to induce ipsiRGC axon outgrowth *in vitro* (**Figure 5A&B**).

**Figure 5.**
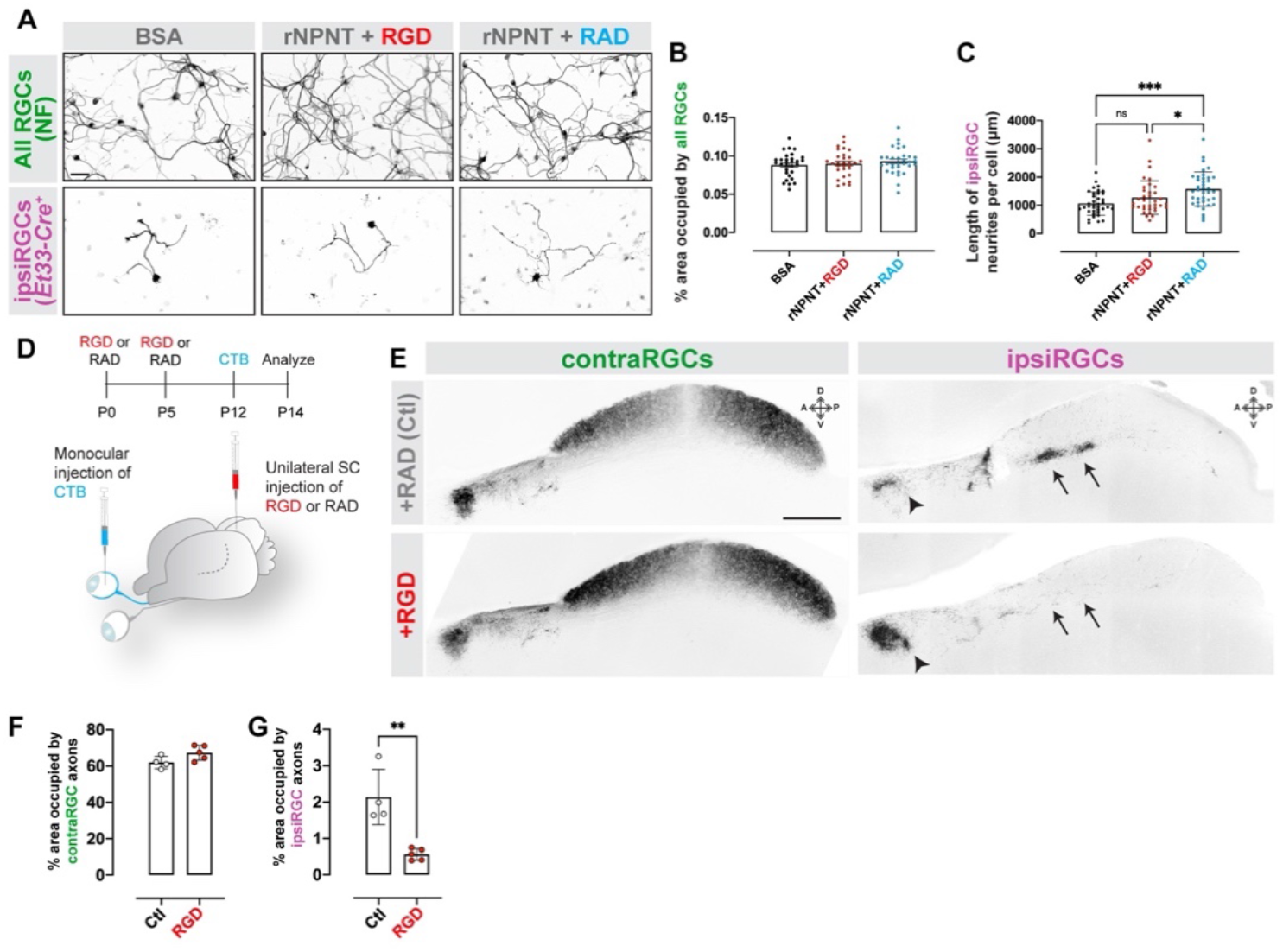
RGC-dependent integrins are required for ipsiRGC innervation of SC. **A.** RGCs *Et33-Cre::Rosa-Stop-tdT* mice were cultured on rNPNT in the presence of integrin-blocking RGD peptides or controls (RAD). All RGCs were labeled by IHC for neurofilament (NF). ipsiRGCs were identified by tdT expression. **B**. Neurite outgrowth of tdT^+^ ipsiRGCs from **F** was quantified by measuring the total length of tdT^+^ neurites. Bars represent mean +/− SD. Data points represent neurites from a single tdT^+^ ipsiRGC from a total of three experiments. P< 0.001 by ANOVA (n=30 cells). **C**. Schematic representation of the timeline and approach for intracollicular delivery of function blocking integrin peptides and intraocular delivery of CTB. **D**. CTB-labeled contra- and ipsiRGCs projections in P14 SC that received control peptides (RAD) or integrin-blocking peptides (RGD). Images depict sagittal sections of SC. Arrows highlight sublamina of SC targeted by ipsiRGC axons which are largely absent from RGD treated SC. Arrowheads highlight ipsiRGC axon targeting of pretectal nuclei in control (RAD) and RGD treated mice. **E,F.** Quantification of the area of SC occupied by contra- (**C**) and ipsiRGC (**D**) projections in. Bars represent means +/− SD. ** indicate P < 0.005 by Student’s t-test (n=4 mice in control group, 5 mice in RGD group). Scale bar in A = 50 μm and in D = 250 μm.

Based on the necessity of RGD-dependent integrins for NPNT to promote ipsiRGC axon outgrowth (**Figure 3**), we tested whether integrins were required for ipsiRGC innervation of SC *in vivo*. Function blocking RGD peptides (or control peptides) were injected into the neonatal SC as ipsiRGC axons were invading SC (**Figure 5C**). Subsequently, RGC projections from both eyes were labeled with CTB. Neonatal intracollicular delivery of RGD peptides led to ipsiRGC axon targeting deficits similar to those observed in *Npnt^fl/fl^::Nes-Cre* and *Npnt^fl/fl^::Vglut2-Cre* mutants (**Figure 5D**). Delivery of RGD peptides resulted in a dramatic loss of ipsiRGC projections in SC, but had little to no impact on contraRGCs projections or ipsiRGCs projections to other retinorecipient regions, such as pretectum (**Figure 5D-F**).

These *in vitro* and *in vivo* data both implicate RGD-dependent integrins in ipsiRGC axon targeting to SC, but which RGC-derived integrins serve as the receptor for NPNT remained unclear. Several transcriptional profiling studies identified distinct integrin subunits expressed in ipsiRGCs (compared to contraRGCs) (2, 25). Thus, we explored whether developing ipsiRGCs generate the β_1_ integrin subunit - an integrin subunit present in the most well-studied NPNT binding integrin (α8β_1_ integrins; (31, 39)) and previously shown to play an essential role in retinotectal targeting in Xenopus (40). *In situ* hybridization revealed a regionally restricted expression of *Itgb1* (the gene encoding the β_1_ integrin subunit) in the ganglion cell layer of the developing mouse retina (**Figure 6A&B**). *Itgb1* expression colocalized with retrogradely-labeled ipsiRGCs in the ventrotemporal crescent of retina (**Figure S5**). To test whether ipsiRGCs generate *Itgb1*, we performed *in situ* hybridization in retinas from *Et33-Cre* mice. We observed expression of *Itgb1* in tdT^+^ ipsiRGCs (**Figure 6C**). *Itga8* mRNA (which encodes α8 integrin) was also generated by tdT^+^ ipsiRGCs in the early postnatal retina of *Et33-Cre::Rosa-Stop-tdT* mice (**Figure 6C&D**). Thus, developing ipsiRGCs generate the NPNT receptor.

**Figure 6.**
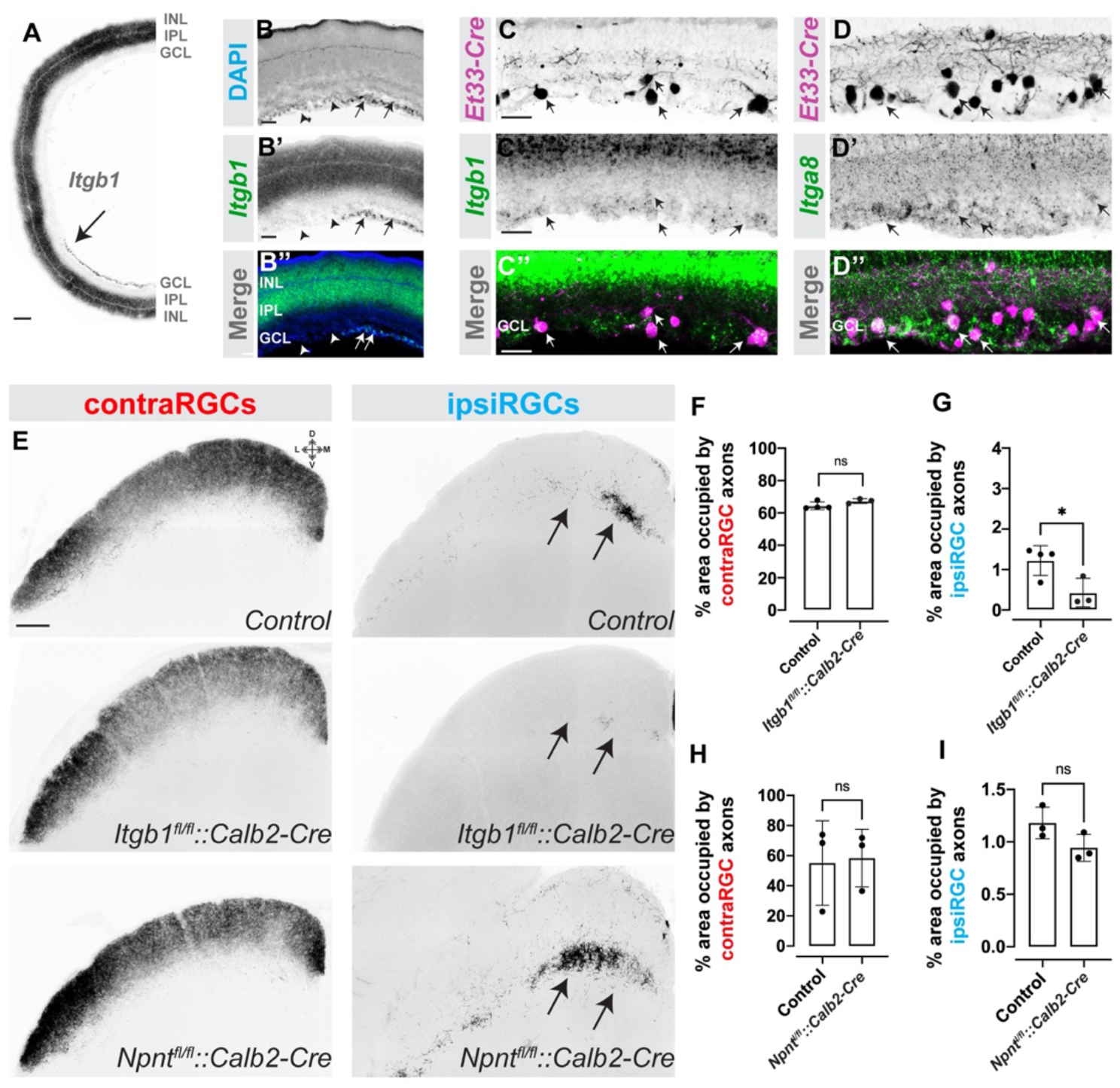
β1 containing integrins are generated by ipsiRGCs and are required for ipsiRGC innervation of SC. **A,B**. ISH for *Itgb1* mRNA in retinal cross sections from P14 wildtype mice. **A** depicts a low magnification image representing almost half of the retina; **B** depicts a high magnification image of *Itgb1* mRNA and DAPI-labeled nuclei. Arrows highlight *Itgb1* expression in regionally-restricted domains of the ganglion cell layer of retina. Arrowheads depict regions of the GCL without appreciable *Itgb1* expression. **C,D**. ISH for *Itgb1* (**F**) and *Itga8* (**G**) mRNAs in retinal cross sections from P12 *Et33-Cre::Rosa-Stop-tdT* mice. Arrows highlight tdT^+^ ipsiRGCs expressing these integrin subunits. **E.** CTB labeling of contra- and ipsiRGCs projections to SC in P14 control, *Itgb1^fl/fl^::Calb2-Cre*, and *Npnt^fl/fl^::Calb2-Cre* mice. Images depict coronal sections of SC. Arrows highlight laminar targeting of ipsiRGC axons in control mice and there absence in *Itgb1^fl/fl^::Calb2-Cre* mice. **F,G**. Quantification of the area of SC occupied by contra- (**F**) and ipsiRGC (**G**) projections in *Itgb1^fl/fl^::Calb2-Cre* in **E**. Bars represent means +/− SD. ns indicates no significance and * indicates P<0.05 by Student’s t-test (n=4 mice in control group and 3 mice in mutant group). **H,I.** Quantification of the area of SC occupied by contra- (**F**) and ipsiRGC (**G**) projections in *Npnt^fl/fl^::Calb2-Cre* in **E**. Bars represent means +/− SD. ns indicates no significance by Student’s t-test (n=3 mice in control group and 3 mice in mutant group). Scale bar in **A** = 200 μm; in **B** = 50 μm; in **C** = 50 μm for **C,D**; in **E** = 200 μm.

Finally, to test whether RGC-derived β_1_ integrin is necessary for ipsiRGC axon targeting of SC, we crossed mice with a conditional allele of *Itgb1 (Itgb1^fl/fl^*) to *Calb2-Cre* mice. Anterograde labeling with CTB was used to assess ipsiRGC and contraRGC projections to SC. While contraRGC projections in *Itgb1^fl/fl^::Calb2-Cre* resembled those in littermate controls, a dramatic loss of ipsiRGC projections was observed in mutants (**Figure 6E-G**). As described above, conditional deletion of NPNT from this same population of Cre-expressing RGCs in *Calb2-Cre* mice did not lead to altered ipsiRGC targeting of SC (**Figure 6E,H&I**). Therefore, taken together, the expression of α8β_1_ integrin by ipsiRGCs and the impairment of ipsiRGC axon targeting by pharmacologically and genetically blocking integrin signaling, suggest that RGC-derived integrins are the receptors that recognize spatially restricted SC-derived NPNT and regulate the assembly of eye-specific, segregated visual pathways.

### Loss of ipsiRGC input to SC disrupts binocular function and impairs innate visual behaviors

Although it has long been appreciated that ipsiRGCs and contraRGCs innervate adjacent, non-overlapping sublaminae of SC, how ipsiRGCs contribute to visual function and behavior is unclear. We sought to answer this question by performing multi-channel extracellular recording from neurons in the anterior SC which receives direct retinal input from both eyes (**Figure 1A**). Briefly, anesthetized mice (*Npnt^fl/fl^::Vglut2-Cre* or controls) were exposed to drifting grating visual stimuli in three ocular conditions: to both eyes simultaneously or with either the ipsilateral or contralateral eye covered (**Figure 7A**). In controls, as we recently reported (41), approximately one-third of SC neurons were driven monocularly (with ~64% of these being driven by the contralateral eye). The remaining SC neurons were binocularly driven and could be divided into four distinct response types: those neurons that respond to stimuli present to either eye or both eyes together (termed simple binocular units [BN-S]), those that respond only when stimuli were presented simultaneously to both eyes (termed emergent binocular units [BN-E]) and those that respond when one eye is covered but not when visual stimuli are presented to both eyes simultaneously (termed binocular units inhibited by the ipsilateral or contralateral eye [BN-IbI or BN-IbC, respectively]) (41)(**Figure 7B and S6**). Recording in *Npnt^fl/fl^::Vglut2-Cre* mutants revealed two important differences. First, we failed to detect neurons that were monocularly driven by the ipsilateral eye (**Figure 7C**). Second, we observed an increase in the proportion of neurons driven monocularly by the contralateral eye and therefore a decrease in the proportion of binocularly driven neurons (**Figure 7B**). When we examined the tuning properties of the units present in *Npnt^fl/fl^::Vglut2-Cre* mutants we did not observe significant differences in orientation selectivity, tuning widths, or linearity of cells in the *Npnt^fl/fl^::Vglut2-Cre* mutant SC (**Figure S6**). Taken together, the absence of neurons driven monocularly by the ipsilateral eye and the decrease in the proportion of neurons that are binocularly responsive in the *Npnt*-deficient mutants are in line with our expectation based on the dramatic loss of ipsiRGC projections to SC in these mutants.

**Figure 7.**
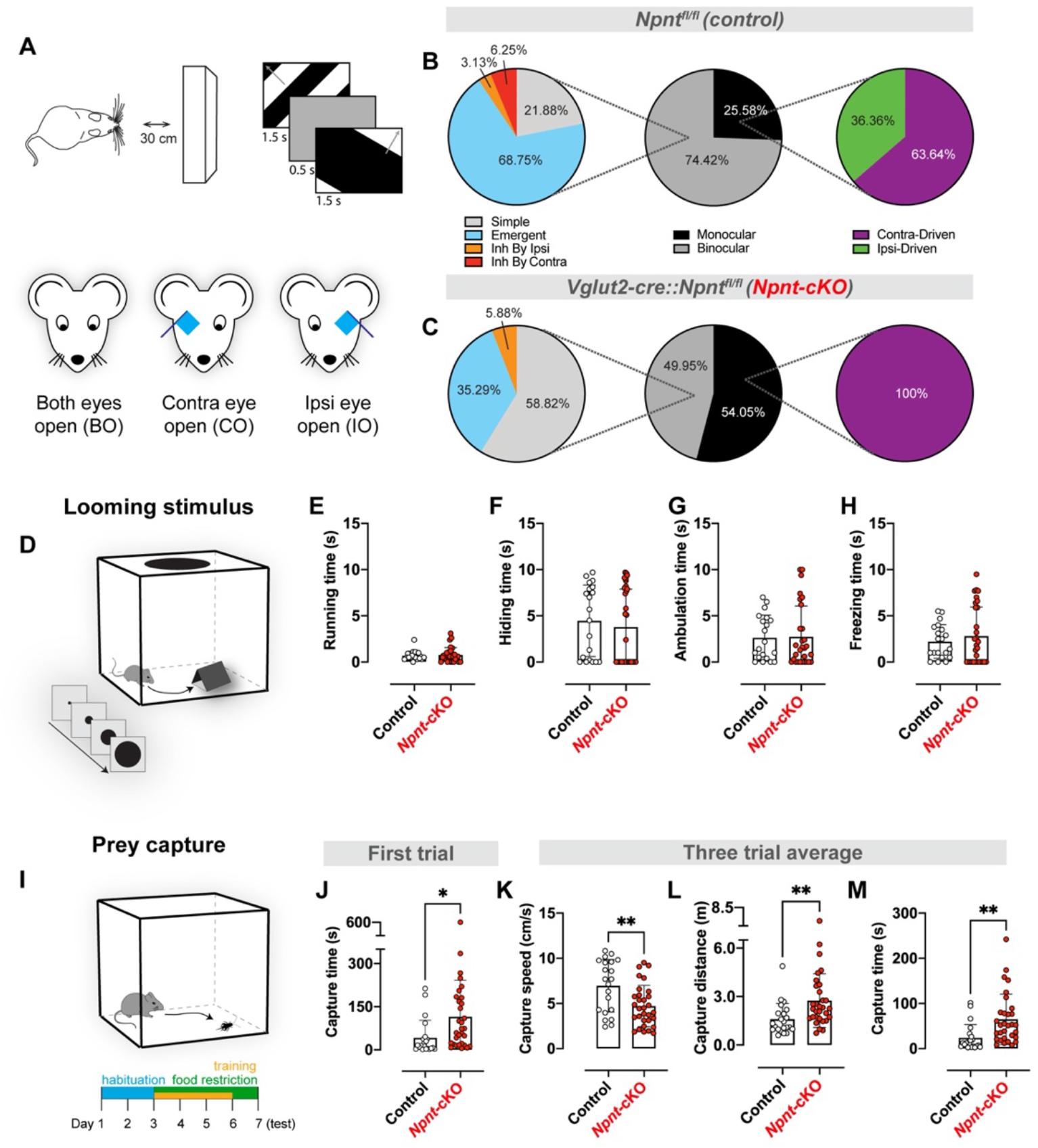
ipsiRGC projections are necessary for binocular neurons and innate visual behaviors. **A.** Schematic representation of visual stimulus paradigm in which drifting gratings were presented directly in front of mice under three ocularity conditions. **B,C.** Proportions of monocular and binocular subtypes of visual neurons identified in the superior colliculi of control (**B**) and *Npnt-cKO* (**C**) mice. **D.** Schematic representation of looming stimulus behavior assay in which an expanding dark disc is presented above the mouse in an enclosed box. **E-H**. Quantification of different aspects of looming stimulus response behavior observed. Specifically, no significant differences (Student’s t-test) were found between control and *Npnt-cKO* mice in running time (**E**), hiding time spent in the shelter (**F**), ambulation time (**G**), and freezing time (**H**). Bars represent means +/− SD, dots represent individual mice. **I.** Schematic representation and timeline of prey capture behavior assay in which mice hunt crickets in an enclosed box. **J-M.** Quantification of different aspects of prey capture behavior observed, specifically comparing control and *Npnt-cKO* mice in capture time during their first trial (**J**), then comparing capture speed (**K**), capture distance (**L**), and capture time (**M**) averaged across three trials. * indicates P<0.05 and ** indicates P<0.005 (Student’s t-test). Bars represent means +/− SD, dots represent individual mice.

Are direct inputs from the ipsilateral retina and binocularly responsive SC neurons important for ethologically relevant visual behaviors? Our anatomical and function studies in *Npnt^fl/fl^::Vglut2-Cre* mutants led us to ask what role direct ipsiRGC innervation may have on SC-related innate visual behaviors. To test this, we assessed the performance of *Npnt^fl/fl^::Vglut2-Cre* mutants (and littermate controls) in two well-established visually guided behavioral tasks: response to a looming spot (42, 43) and the prey capture task (44, 45) (**Figure 7D&E**). Mutants lacking NPNT (and therefore ipsiRGC projections to SC) performed similar to controls in terms of freezing time, running time, hiding time, or ambulation time when presented with a dark looming stimulus (**Figure 7E-H**). In contrast, the ability of these mutants to capture prey was significantly impaired (**Figure 7J-M**). Compared to littermate controls, *Npnt^fl/fl^::Vglut2-Cre* mutants took longer to capture prey, travelled farther during prey pursuit, and moved slower during prey pursuit (**Figure 7J-M**). These data suggest that direct inputs from the ipsilateral retina are critical for some, but not all, SC-mediated visual behaviors.

## Discussion

Establishing precise and stereotyped cell type-specific circuits is a major challenge during brain development. Due to its accessibility, the subcortical visual system has served as a model for understanding mechanisms that underlie fundamental aspects of circuit formation: axon outgrowth and guidance, target selection, synaptogenesis and activity-dependent refinement. Despite considerable advances in our understanding of many of these processes, what has been lacking is a mechanism that drives individual subtypes of RGCs to project axons to regionally-restricted domains of retinorecipient nuclei which serves to parse different types of visual information into anatomically distinct parallel channels. Here, we identify a specific molecular matching mechanism that drives laminar targeting of ipsiRGC axons and establishes an eye-specific, segregated parallel visual pathway.

### Cell-ECM molecular matching mechanisms underlie laminar targeting of RGC axons

The assembly of cell type-specific circuits between the retina and brain relies on a combination of intrinsic transcriptional codes, cell-cell/cell-ECM interactions, and activity-dependent mechanisms (9, 10, 46–49). Focusing on ipsiRGCs, these steps include the guidance of axons from the ventrotemporal crescent of retina to the optic disc, the divergence of these axons from contraRGC at the optic tract, the selection of retinorecipient targets, the generation of eye-specific domains (through activity dependent refinement or through laminar targeting depending on the brain region) and the formation, maturation, and refinement of synaptic connections (46, 50–53). Intrinsic transcriptional codes not only differentiate ipsiRGCs from contraRGCs (2, 25), they also provide these ipsilateral projecting cells with a unique repertoire of cell surface receptors to respond uniquely to molecule cues as they course into the brain. Indeed, cell-cell and cell-ECM mechanisms drive the divergence of ipsiRGC axons at the optic chiasm and their homophilic fasciculation in the optic tract (51, 52, 54, 55).

Here, we add to this rich literature by revealing that a novel cell-ECM mechanism drives laminar targeting of ipsiRGC axons in SC. NPNT, a regionally restricted ECM glycoprotein containing epidermal growth factor like-repeats, integrin-binding motifs, and a MAM (meprin-A5 protein-receptor protein phosphatase) domain (31, 32, 39), is generated in the SC sub-lamina innervated by ipsiRGCs. The protein domains of NPNT classify it as a member of an EGF-like family of adhesive glycoproteins (including laminins, thrombospondins, and tenascins). Members of this family of glycoproteins have well-documented roles in tissue development and morphogenesis, including in promoting axonal outgrowth (56, 57). Outside of the brain, NPNT plays critical roles in renal development (31, 36). Studies here now demonstrate novel roles for this adhesive glycoprotein in the brain, in that NPNT is sufficient to promote ipsiRGC neurite growth *in vitro* and is necessary for ipsiRGC innervation of SC *in vivo*. Thus, like has been shown for other vertebrate visual systems (58–60), an ECM glyco-code specifies precise subtype-specific laminar targeting in the mammalian retinocollicular circuit. Moreover, these results redefine the biological role of NPNT as a brain-derived laminar targeting cue.

Intrinsic differences in ipsiRGCs allow them to uniquely respond to SC-derived NPNT. A number of cell surface receptors, including integrins, are enriched in ipsiRGCs compared to other subtypes of RGCs (2, 25, 26). We now show that developing ipsiRGCs express α8β_1_ integrin. This integrin heterodimer was initially identified as the NPNT receptor (31) and was shown to promote axon outgrowth *in vitro* (61). In support of a role for this receptor in retinocollicular targeting in mice, previous studies showed that blocking β_1_ containing integrins disrupts retinotectal circuit formation in Xenopus (40). Here we show that disrupting integrin RGD-binding with function blocking peptides impairs ipsiRGC axon growth on rNPNT *in vitro* and disrupts ipsiRGC innervation of SC *in vivo*. Genetic loss of β_1_ integrins from RGCs likewise results in a loss of ipsiRGC innervation of rodent SC. Thus, integrins are necessary receptors for the laminar targeting of ipsiRGC axons in mice. This is an important difference from studies in zebrafish which identified roles for two integrin-binding ECM molecules -- CollagenIVα5 and Reelin -- in retinotectal targeting (58, 60). Although both CollagenIVα5 and Reelin are capable of signaling through β_1_ integrins (although not the α8β_1_ integrin; (62, 63)), β_1_ integrins are not required for the laminar targeting of RGCs in fish (58). While these differences could reflect differences in eye-specific projection patterns in these species, they may simply highlight the numerous molecular matching mechanisms necessary to pair axons from the single RGC subtypes to the correct target lamina or cells in the brain.

This raises the question of whether a unique receptor-ligand recognition mechanism is required for the laminar targeting of each RGC subtype. It is certainly possible. Loss of SC-derived NPNT impairs ipsiRGC projections to SC but not those of the On-Off direction-selective RGCs labeled in *Trhr-GFP* mice (**Figure 4**; (15)). It is also possible that a gradient of ECM molecules (such as NPNT) could be used to pattern the targeting of axons from multiple subtypes of RGCs in the mammalian SC. This is the case in zebrafish tectum where opposing gradients of attractive reelin and repellent Slit2 are thought to convey positional information for invading RGC axons (59, 60). Such a mechanism would greatly reduce the number of receptorligand recognition mechanisms to pattern laminar targeting (and similar overlapping gradients drive topographic map formation in mammalian retinorecipient zones; (16, 47). Here, we show that a large subset of ipsiRGCs are αRGCs based on the expression of *Spp1* mRNA or based on their immunoreactivity for SMI-32 (**Figure 1E&F**) (2, 64, 65). Contralateral projecting αRGCs target SC sublamina that are just dorsal to the strata innervated by ipsiRGCs (19), so it is possible that they share some of the machinery to respond to NPNT. While a shared mechanism may drive αRGC lamina targeting in SC, the same mechanism may not be responsible for cell type-specific circuits in other retinorecipient zones. As an example, NPNT is necessary for ipsiRGC innervation of SC, but not in visual thalamus, suggesting unique cell-ECM recognitions mechanisms are required in each central target. Similar nuclei-specific difference in targeting mechanisms have been demonstrated for intrinsically photosensitive RGCs (ipRGCS) which require the ECM molecule Reelin to innervate thalamic nuclei but not the suprachiasmatic nucleus (66, 67). Regardless of whether each synaptic lamina in SC contains unique recognition molecules or whether gradients of cues diffuse across mouse SC, it is clear that additional cell-ECM and cell-cell recognition mechanisms remain to be elucidated for a complete understanding of how cell type-specific visual circuits form in the developing mouse brain.

### Multiple pathways convey information from the ipsilateral eye to the SC

The SC is responsible for driving goal-directed eye-movements and an emerging number of innate visual behaviors (42–45, 68, 69). Here we asked whether direct input from the ipsilateral eye was required for some of these innate behaviours. Information from the ipsilateral eye would likely generate binocular responses in SC and could contribute to depth perception or to saccade-like eye movements to place objects of interest into central areas of the visual field for enhanced feature detection (12, 70) but see (71). Perhaps it is not surprising then that *Npnt*-deficient mutants exhibit defects in prey capture, an ethologically relevant behaviour that requires shifts in gaze to stabilize the visual scene as objects are tracked (72). To our knowledge, this is the first evidence for a behavioral role of direct connections between the SC and ipsilateral eye. In contrast, behaviors that do not require the same precise orientations of the visual field (such as responses to looming stimulus) do not appear to require ipsiRGC innervation of SC.

While these analyses identified key roles for ipsiRGC innervation of SC in an innate behavior, functional analysis of cells in the SC of *Npnt*-deficient mice unexpectedly informed us about the circuits underlying binocularly responsive neurons. While such neurons have not been well-studied in mammals with laterally oriented eyes (such as rodents), they have been in higher mammals with forward facing eyes, given the role of the SC in orienting head and eye movement in these species (73–75). It was for this reason we were surprised when we discovered that a substantial proportion of cells in the anterior mouse SC were binocularly responsive (41). Four distinct subtypes of binocularly responsive cells were identified (41). Here, we sought to explore how the responses of these cells might change in the *Npnt*-deficient mice which lack direct innervation by ipsiRGCs. Not surprisingly, we found no cells in *Npnt^fl/fl^::Vglut2* mutants whose responses were driven only by the ipsilateral eye (**Figure 7**). We also found a significant decrease in the proportion of binocularly responsive neurons in these mutants. However, a large number of binocularly responsive neurons remained in these mutants despite the apparent lack of ipsiRGC axons. Of course, it is possible that these responses are driven by the few ipsiRGC fibers that do remain in the SC of these mutants. However, in our opinion, the paucity of these fibers makes this seem unlikely. Instead, we interpret these results to suggest that indirect pathways may exist that transmit information from the ipsilateral eye to the SC. Such indirect pathways have been identified in post-metamorphic frogs which lack direct ipsiRGC-SC circuits (76). Thus, taken together, evidence from our anatomical, functional, and behavior studies in *Npnt*-deficient mice highlight the necessity of parsing visual information into parallel visual streams for some behaviors, but also highlight that sensory systems are more complex than simple parallel pathways.

## Materials and Methods

A complete description of materials and methods is available in the supplemental materials.

### Mouse Lines and Husbandry

C57BL/6 mice were obtained from Charles River Laboratories (Wilmington, MA, USA). *Pvalb-Cre, Nes-Cre, Gad2-Cre, Calb2-cre, Vglut2-cre, Sst-Cre, Rosa-stop-tdT*, and *Thy1-stop-YFP* mice were obtained from Jackson Labs (stock # 008069, 003771, 010802, 010774, 016963, 013044, 007909). *Trhr-EGFP* mice (stock # 030036-UCD) were obtained from MMRRC. Conditional allele of *Npnt (Npnt^fl/fl^*) mice were kindly from Dr. Denise K. Marciano (University of Texas Southwestern)(77). Conditional allele of *Itgb1 (Itgb1^fl/fl^*) and *Aldh1l1-EGFP* mice were provided by Dr. Stefanie Robel (Virginia Tech)(78, 79). Et33-Cre mice were kindly from Dr. Colenso Speer (UM) (26). Mice were housed in a 12 hr dark/light cycle and had ad libitum access to food and water. All experiments were performed in compliance with National Institutes of Health (NIH) guidelines and protocols and were approved by the Virginia Polytechnic Institute and State University Institutional Animal Care and Use Committee (IACUC).

### Tissue preparation and immunohistochemistry (IHC)

Fluorescent IHC was performed on 20-μm cryosectioned PFA-fixed brain tissue (66, 80–83). Tissue slides were allowed to air-dry for 15 min before being incubated with blocking buffer (2.5% normal goat serum, 2.5% BSA, and 0.1% Triton X-100 in PBS) for 30 min. Primary antibodies were diluted in blocking buffer and incubated on tissue sections overnight at 4°C. The following antibodies and dilutions were used: mouse anti-Brn3a (diluted 1:125, Millipore), rabbit anti-RFP (diluted 1:500, Rockland), rabbit anti-Opn4 (diluted 1:2000, Dr. C.K. Chen’s lab (66)), mouse anti-SMI32 (diluted 1:1000, Covance), rabbit anti-RBPMS (diluted 1:500, PhosphoSolutions), rabbit anti-GFP (diluted 1:250, Invitrogen), mouse anti-NeuN (diluted 1:200, Millipore), rabbit anti-GFAP (1:1000, DAkoCytomation), rabbit anti-Iba1 (1:500, Wako), mouse anti-GAD67 (diluted 1:500, Millipore), rabbit anti-calbindin (diluted 1:2500, Swant), rabbit anti-calretinin (diluted 1:2000, Swant), mouse anti-synaptophysin (diluted 1:500, SySy), goat anti-nephronectin (diluted 1:40, R&D systems). On the next day, tissue slides were washed in PBS, and secondary antibodies diluted 1:1,000 in blocking buffer were applied to slides for 1 hr at RT. After thorough washes in PBS, tissue slides were coverslipped with VectaShield (Vector Laboratories). Images of tissue were acquired on a confocal microscope (LSM 700; Zeiss) equipped with a 20x air Plan-Apochromat objective (NA 0.8; Zeiss) and a 40Å~ oil EC Plan-NeoFluor objective (NA 1.3; Zeiss). When comparing different ages of tissues or between genotypes, images were acquired with identical parameters, and similar gamma adjustments were made to age-matched mutant and control images in Adobe Photoshop or ImageJ. A minimum of three animals (per genotype and per age) were compared in all IHC experiments.

### Riboprobe making and *in situ* hybridization

*In situ* hybridization (ISH) was performed on 20-μm sagittal or coronal cryosectioned tissues (Su et al., 2010, 2016, 2020). Antisense riboprobes were generated from full-length *Npnt* (MMM1013-202708550), *Syt1* (MM1013-9199901), *Itgb1* (MMM1013-202859073) and *Itga8* (MMM1013-202705925) Image Clones (Dharmacon) as described previously (82–84). Antisense riboprobes were generated against a 599-bp fragment of *Sst* (corresponding to nt 1-599 of NM_009215.1), a 973bp fragment of *Spp1* (Corresponding to nt 309-1279 of NM_001204201.1), a 625bp fragment of *Gda* (Corresponding to nt 1884-2508 of NM_010266.1), a 580bp fragment of *Vglut2* (Corresponding to nt 2190-2769 of NM_080853.2) and a 982bp fragment of *Gad1* (Corresponding to nt 1015-1996 of NM_008077.2) were PCR-cloned into pGEM Easy T vector (Promega).

In brief, riboprobes were synthesized using digoxigenin (Dig) or fluorescein (Fl)-labeled UTP (Roche) and the MAXIscript In Vitro Transcription kit (Ambion). Probes were hydrolyzed to 400-500 nt. Tissue sections were fixed in 4% PFA for 10 min, washed with DEPC-PBS three times, and incubated in proteinase K solution for 10 min. Subsequently, slides were washed with DEPC-PBS, fixed with 4% PFA for 5 min, washed with DEPC-PBS, and incubated in acetylation buffer for 10 min. Slides were then permeabilized in 1% Triton X-100 for 30 min and washed with DEPC-PBS. Endogenous peroxidase was blocked by incubation in 0.3% H2O2 for 30 min. Tissue sections were equilibrated in hybridization buffer for 1 h and incubated with probes at 65°C overnight. After washing in 0.2x SSC at 65°C, bound riboprobes were detected by horseradish peroxidase (POD)-conjugated anti-Dig (1:2000, Roche) or anti-Fl (1:2000, Roche) antibodies followed by fluorescent staining with Tyramide Signal Amplification system (PerkinElmer). After mounting sections in VectaShield, images were obtained on a Zeiss LSM 700 confocal microscope equipped with a 20x air Plan-Apochromat objective (NA 0.8). A minimum of three animals per genotype and age were compared in ISH experiments.

### Quantitative real-time PCR

RNA was isolated using the BioRad Total RNA Extraction from Fibrous and Fatty Tissue kit (BioRad). cDNAs were generated from 500ng RNA with the Superscript II Reverse Transcription First Strand cDNA Synthesis kit (Invitrogen). Quantitative real-time PCR (qPCR) was performed on a Chromo 4 Four Color Real-Time system (BioRad) using iTaq SYBRGreen Supermix (BioRad; Su et al., 2016).*Npnt* primers for qPCR were 5’- GAG CTG GCA CCT TAT CCT TG-3’ and 5’- CTA TGG AGG CAG GAT TGA CTG −3’. *Itga8* primers for qPCR were 5’- TTG TGA GCT CTC ACT GTG GC −3’ and 5’- AGA TAC CGT TTG ACA CCA CCA −3’. *Itgb1* primers for qPCR were 5’- GCA ACGCAT ATC TGG AAA CTT G −3’ and 5’- CAA AGT GAA ACC CAG CAT CC −3’. *18s* primers for qPCR were 5’- GGA CCA GAG CGA AAG CAT TTG −3’ and 5’- GCC AGT CGG CAT CGT TTA TG −3’. qPCR primers were designed over introns. The following cycling conditions were used with 10 ng RNA: 95°C for 30 s, followed by 40 cycles of amplification (95°C for 5 s, 60°C for 30 s, 55°C for 60 s, read plate) and a melting curve analysis. Relative quantities of RNA were determined using the ΔΔ-CT method. A minimum of double experiments (each in triplicate) was run for each gene. Each individual run included separate *Gapdh* control reactions.

### Intraocular injections of anterograde tracers

Intraocular injection of cholera toxin subunit B (CTB) conjugated to Alexa Fluor 488 or Alexa Fluor 555 (Invitrogen) was performed as described previously (66, 67). Briefly, mice were anesthetized with hypothermia (<P7) or by isoflurane vapors (>P7). The sclera was pierced with a sharp-tipped glass pipette and excess vitreous was drained. Another pipette, filled with a 1ug/ul solution of CTB, was inserted into the hole made by the first pipette. The pipette containing the CTB was attached to a picospritzer and a prescribed volume (1 to 2 μl at P3 to P10 and 2 to 3 μl for ages >P10) of solution was injected into the eye. After 1 to 2 days, mice were killed and brains were fixed in 4% paraformaldehyde. A total of 100 μm sagittal or coronal sections were sectioned on a vibratome (Microm HM 650 V; Thermo Scientific, Waltham, MA, USA) and mounted in Vectashield Mounting medium (Vector lab). Retinal projections were analyzed from at least 3-5 animals for each age and genotype. To quantify the spatial extent of superior colliculus (SC) or lateral geniculate nucleus (LGN) innervation by retinal axons, serial sagittal or coronal sections encompassing the entire SC or LGN mounted on slides. Images were acquired on a Zeiss LSM 700 confocal microscope. Use ImageJ to quantify the percentage of area occupied by retino-superior colliculus or retinogeniculate targeting signals.

### Delivery of peptides or retrograde tracers into SC

RGD or RAD peptide injection was performed as follows. Postnatal zero (P0) mice were anesthetized with hypothermia, and the skin overlying the SC was opened and reflected. A sharp-tipped glass pipette filled with 1μL of 100μM RGD or RAD solution was then inserted through the thin skull and into the anterior neonatal superior colliculus. Solution within the pipette (i.e. RGD or RAD) was slowly depressed into the SC pneumatically, via a picospritzer. After 5 days (P5), mice were anesthetized a second time and were injected with the same volume, solution, and SC location. After another 7 days (P12), mice received monocular injections of Alexa Fluor 555-conjugated CTB as described previously. After 2 days, mice were euthanized and brains were fixed in 4% paraformaldehyde. RGC projections were analyzed in 100μm sagittal sections that were cut on a vibratome and mounted as described above.

To retrogradely label ipsiRGCs, intracollicular injections of CTB were performed. A similar procedure as described above was applied to P3 mice. 1μL of 1μg/mL Alexa Fluor 555-conjugated CTB was injected into the anterior SC via a picospritzer. Three days after intracollicular delivery, mice were euthanized, fixed with 4% PFA, and retinas removed.

### AAV virus injection

Viral tracing was done as described in (34). Briefly, AAV2/1-hSyn-Cre-WPRE-hGH (2.5 × 10^13^ GC/mL, here referred to as AAV1-Cre) was used to monosynaptically label retinorecipient neurons in the SC. Briefly, mice were anesthetized with isoflurane and 1.2 μl of AAV-Cre virus was monocularly injected at an approximate 45° angle relative to the optic axis. AAV1-Cre was a gift from James M. Wilson (Addgene viral prep #105553-AAV1; RRID:Addgene_105553). Animals were sacrificed and perfused with PFA as described above 6-10 weeks after injection.

### RGC immunopanning

We referenced the protocol outlined in (85) in developing our approach to purifying and culturing RGCs. Retinas of *Et33-Cre::tdT* (~P5) were dissected using a dissection microscope and dissociated using papain. We immunopanned RGCs from retinal suspension using Thy1.2 antibody (CD90, 1:800, Bio-RAD) and 0.02% BSA (Sigma-Aldrich) for at least 2 hours at room temperature (positive panning dish). Two 15-cm petri dish were incubated with BSL-1 (5μg/mL, Vector Labs) in D-PBS for at least 2 hours at room temperature (negative panning dishes). RGCs were detached from the dish by pipetting prewarmed RGC growth medium directly to the dish. Cells were collected by centrifuging the tube at 200xg for 12 minutes at 25°C. ~70,000 cells/well were seeded in a 8-chamber slide coated with either 10μg/mL rNPNT (R&D system), 2μg/mL BSA, or 1X Poly-D-Lysine (Sigma-Aldrich). Chamber-slide wells were coated with rNPNT or BSA for 2 hours at 37°C before seeding cells. Medium was changed every other day. For peptide treatment, medium was changed on the second day with RGC growth medium containing either 10μM GRGDSP or 10μM GRADSP. After 5 days, cells were fixed for immunocytochemistry. Please see SI Appendix for more details.

### Electrophysiology

Visual response properties were determined as previously described (86), with minor modifications. Briefly, isoflurane-anesthetized adult mice were head-fixed and a 16-channel silicon multi-electrode (Neuronexus Technologies) was inserted into at an angle of 45 deg to the midline and 45 deg to the horizontal plane through a craniotomy located ~1.5 mm lateral to the midline and ~1.5 anterior to lambda. Electrodes were labeled with 1,1’-Dioctadecyl-3,3,3’,3’-Tetramethylindocarbocyanine Perchlorate (Invitrogen) and localization was confirmed *post hoc* via fluorescent microscopy. Multiunit signals were acquired at ~25 kHz and filtered between 0.7-7 kHz using a System3 Workstation (Tucker-Davis Technologies). Individual units were identified *post hoc* using independent components analysis. Visual stimuli were displayed on an LCD monitor subtending ~80 x 50 deg of visual space and placed directly in front of the animal. Stimuli consisted of drifting square waves presented at 12 different orientations and 6 different spatial frequencies, each presented 5-7 times. In addition, full-field flash and gray screens were presented to provide robust visual stimulus and determine spontaneous firing rate, respectively. Stimuli were presented in three ocular conditions: to both eyes together (both open, BO), with the ipsilateral eye covered (contralateral open, CO), and with the contralateral eye covered (ipsilateral open, IO). Based on their responsiveness under each condition, units were classified into one of seven potential types (41): monocularly-driven by the contralateral eye (BO^+^CO^+^IO^-^), monocularly-driven by the ipsilateral eye (BO^+^CO^-^IO^+^), binocular simple (BO^+^CO^+^IO^+^), binocular emergent (BO^+^CO^-^O^-^), binocular inhibited by ipsilateral (BO^-^CO^+^IO^-^), binocular inhibited by contralateral (BO^-^CO^-^O^+^), or binocular cross-inhibited (BO^-^CO^+^IO^+^).

### Looming assay

Looming stimuli were presented to the animals in white rectangular arena (47×37×30 cm) with an opaque shelter placed in a corner. Entrance of the shelter was facing the center of the arena. The arena was diffusely and evenly illuminated from above and was located within a light-proof and sound isolated room to maintain constant environmental conditions. A camera with frame rate of 30 FPS for capturing mouse’s behavior was secured to the stand next to the arena. All mice were tested only once to avoid a habituation to the looming stimulus. At the beginning of test, animals were let freely investigate the arena and the shelter for the period of 10 mins before the recording started. We started the video capturing approximately 10 sec prior to looming stimulus and the looming stimulus began when the animal was moving around the center of arena. Videos were recorded 10 sec prior, during and after looming stimulus. The animal’s behavior was scored manually during 10 sec of looming stimulus using ANY-maze software. Similar to (87), we scored 4 types of behavior - freezing, running, hiding and ambulation. Freezing was defined as period of one or more seconds in which the animal was completely immobile. Running was scored as activity in which the mouse started to move at least two times faster, then the average speed before stimulus onset. Hiding was defined as activity when the mouse was completely hidden in the shelter. Ambulation was defined as all other locomotor activity performed in the open arena. Please see SI Appendix for more details.

### Prey capture

The task was recorded in a rectangular, white acrylic arena 47 cm long x 37 cm wide x 30 cm high using the ANY-maze software. The arena was diffusely and evenly illuminated from above and was located within a light-proof and sound isolated room to maintain constant environmental conditions. The floor of the arena was cleaned between each trial and mice with 30% EtOH and let to dry out completely. In order to perform this task, we followed the protocol from the study of (45). Each animal in our protocol underwent a 6-day acclimation protocol followed by testing day. During testing, each mouse was given three 10-min trials to catch a cricket within the arena. If the mouse caught the cricket within these 10 min, the trial was scored as a capture success, and the capture time for that trial was recorded. If a cricket was not captured within 10 min, the mouse was removed from the arena for 1 min and returned into the arena with a new cricket to start the next trial. Mean capture time and average speed during the hunt for each mouse on each day was calculated. All tested animals achieved 100% capture success on testing day. Please see SI Appendix for more details.

### Quantification and Statistical Analysis

To quantify retinal axons in SC, pretectum, and LGN, fluorescent signals in confocal micrographs were binarized using ImageJ and the brain region was encircled to measure area and quantify the fraction of that area that is occupied by binarized signals. For RGC immunopanning, fluorescent signals (of either *Et33^+^* or NF^+^ RGC neurites) were binarized in ImageJ and the fraction of the total area of the field of view they occupied was quantified. To measure neurite length in RGC immunopanning, we measured the total length of *Et33^+^* or NF^+^ neurites per field of view in ImageJ before normalizing to each cell. When comparing measurements between mutants and controls, we determined statistical significance either by Student’s t-test or by ANOVA where appropriate (indicated in figure legends) using GraphPad Prism (version 8.0.; RRID:SCR_002798). Differences were considered significant when P < 0.05 and P-values were indicated in figure legends. No data or animals were excluded from any of the analyses.

## Acknowledgments

The authors thank Dr. Anthony LaMantia and colleagues at Virginia Tech for valuable comments on this manuscript. This work was supported in part by the following grants from the National Institutes of Health: EY021222 (M.A.F), EY030568 (M.A.F), NS105141 (M.A.F.), NS113459 (U.S.), and EY029874 (J.W.T.). The authors are grateful to Dr. Denise Marciano (UTSW) for providing *Npnt^fl/fl^*mice, Dr. Stefanie Robel (VT) for providing *Itgb1^fl/fl^* mice, to Dr. J.M. Wilson for providing AAV1-Cre virus, to Colenso Speer (UM) for providing *Et33-Cre* mice, and to Dr. C.K. Chen (BCM) for providing anti-Opn4 antibodies.

## Supplemental Methods

### Mouse Lines and Husbandry

C57BL/6 mice were obtained from Charles River Laboratories (Wilmington, MA, USA). *Pvalb-Cre, Nes-Cre, Gad2-Cre, Calb2-cre, Vglut2-cre, Sst-Cre, Rosa-stop-tdT*, and *Thy1-stop-YFP* mice were obtained from Jackson Labs (stock # 008069, 003771, 010802, 010774, 016963, 013044, 007909). *Trhr-EGFP* mice (stock # 030036-UCD) were obtained from MMRRC. The following primers were used for genotyping Cre-expressing mice: *Cre*-F: 5’- CGT ACT GAC GGT GGG AGA AT −3’; *Cre*-R: 5’- TGC ATG ATC TCC GGT ATT GA −3’. The following primers were used for genotyping GFP- and tdT-expressing mice: *GFP*-F: 5’- AAG TTC ATC TGC ACC ACC G −3’; *GFP*-R: 5’- TCC TTG AAG AAG ATG GTG CG −3’; *tdT*-F: 5’- ACC TGG TGG AGT TCA AGA CCA TCT −3’; *tdT-R:* 5’- TTG ATG ACG GCC ATG TTG TTG TCC −3’. Conditional allele of *Npnt* (*Npnt^fl/fl^*) mice were kindly from Dr. Denise K. Marciano (University of Texas Southwestern)(1). Primers for *Npnt* genotyping are *Npnt*-F: 5’-CAG TCC ATC CTG ATC ACT GGC TGT A-3’; *Npnt*-R: 5’-GCA ACC TTC AGC GTC CC-3’. Conditional allele of *Itgb1 (Itgb1^fl/fl^*) and *Aldh1l1-EGFP* mice were provided by Dr. Stefanie Robel (Virginia Tech)(2, 3). Primers for *Itgb1* genotyping are *Itgb1-F:* 5’- AGG TGC CCT TCC CTC TAG A; *Itgb1-R:* 5’- GTG AAG TAG GTG AAA GGT AAC-3’. Mice were housed in a 12 hr dark/light cycle and had ad libitum access to food and water. All experiments were performed in compliance with National Institutes of Health (NIH) guidelines and protocols and were approved by the Virginia Polytechnic Institute and State University Institutional Animal Care and Use Committee (IACUC).

### Riboprobe making and *in situ* hybridization

*In situ* hybridization (ISH) was performed on 20-μm sagittal or coronal cryosectioned tissues (Su et al., 2010, 2016, 2020). Antisense riboprobes were generated from full-length *Npnt* (MMM1013-202708550), *Syt1* (MM1013-9199901), *Itgb1* (MMM1013-202859073) and *Itga8* (MMM1013-202705925) Image Clones (Dharmacon) as described previously (4–6). Antisense riboprobes were generated against a 599-bp fragment of *Sst* (corresponding to nt 1-599 of NM_009215.1), a 973bp fragment of *Spp1* (Corresponding to nt 309-1279 of NM_001204201.1), a 625bp fragment of *Gda* (Corresponding to nt 1884-2508 of NM_010266.1), a 580bp fragment of *Vglut2* (Corresponding to nt 2190-2769 of NM_080853.2) and a 982bp fragment of *Gad1* (Corresponding to nt 1015-1996 of NM_008077.2) were PCR-cloned into pGEM Easy T vector (Promega) with the following primers: *Sst*: 5’-AGC GGC TGA AGG AGA CGC TAC-3’ and 5’-CGC CAT AAT CTC ACC ATA ATT TTA-3’; *spp1*: 5’-AAT CTC CTT GCG CCA CAG-3’ and 5’-TGG CCG TTT GCA TTT CTT-3’; *Gda*: 5’-GAG AGG GCA CAA GCT AGA CAT T-3’ and 5’-CCA TAA TGC TTT AGG GAC TTG C-3’, *Vglut2:* 5’-CCA AAT CTT ACG GTG CTA CCT C-3’ and 5’-TAG CCA TCT TTC CTG TTC CAC T-3’ and *Gad1:* 5’-TGT GCC CAA ACT GGT CCT-3’ and 5’-TGG CCG ATG ATT CTG GTT-3’.

In brief, riboprobes were synthesized using digoxigenin (Dig) or fluorescein (Fl)-labeled UTP (Roche) and the MAXIscript In Vitro Transcription kit (Ambion). Probes were hydrolyzed to 400-500 nt. Tissue sections were fixed in 4% PFA for 10 min, washed with DEPC-PBS three times, and incubated in proteinase K solution (1 μg/ml proteinase K, 50 mM Tris, pH 7.5, and 5 mM EDTA) for 10 min. Subsequently, slides were washed with DEPC-PBS, fixed with 4% PFA for 5 min, washed with DEPC-PBS, and incubated in acetylation buffer (1.33% triethanolamine, 20 mM HCl, and 0.25% acetic anhydride) for 10 min. Slides were then permeabilized in 1% Triton X-100 for 30 min and washed with DEPC-PBS. Endogenous peroxidase was blocked by incubation in 0.3% H2O2 for 30 min. Tissue sections were equilibrated in hybridization buffer (1X prehybridization, 0.1 mg/ml yeast tRNA, 0.05 mg/ml heparin, and 50% formamide) for 1 h and incubated with probes at 65°C overnight. After washing in 0.2x SSC at 65°C, bound riboprobes were detected by horseradish peroxidase (POD)-conjugated anti-Dig (diluted1:2000, Roche) or anti-Fl (diluted1:2000, Roche) antibodies followed by fluorescent staining with Tyramide Signal Amplification (TSA) systems (PerkinElmer). After mounting sections in VectaShield, images were obtained on a Zeiss LSM 700 confocal microscope equipped with a 20x air Plan-Apochromat objective (NA 0.8). A minimum of three animals per genotype and age were compared in ISH experiments.

### RGC immunopanning

We referenced the protocol outlined in (7) in developing our approach to purifying and culturing RGCs. A 15-cm petri dish was incubated with goat anti-mouse IgG+IgM (H+L) (1:300, Jackson ImmunoResearch) in 50mM Tris-HCl (pH9.5) at 4°C overnight. Then, the dish was washed with D-PBS (Thermofisher) three times and coated with D-PBS containing Thy1.2 antibody (CD90, 1:800, Bio-RAD) and 0.02% BSA (Sigma-Aldrich) for at least 2 hours at room temperature (positive panning dish). Two 15-cm petri dish were incubated with BSL-1 (5μg/mL, Vector Labs) in D-PBS for at least 2 hours at room temperature (negative panning dishes). Retinas of *Et33-Cre::tdT* (~P5) were dissected using a dissection microscope. Retina Tissues were dissociated at 35°C water bath for 20 minutes by 165 units of Papain (Worthington Biochemical) in 10mL of D-PBS (ThermoFisher). 2mg of L-cycteine (Sigma-Aldrich) and 250 units of DNase (Worthington Biochemcial) were added in papain solution to activate papain and degrade DNA. the pH of papain solution was neutralized by 1N NaOH. After 20 minutes, dissociated retina tissues were collected by centrifuging the tube at 200xg for 3 minutes at 25°C. To isolate single cell suspension and to stop papain reaction, the tissue pellet was triturated with low-Ovomucoid solution (pH 7), containing 1.5 mg/mL of Trypsin inhibitor (Worthington Biochemical) and 1.5 mg/mL BSA (Sigma-Aldrich) in D-PBS, by pipetting up and down three to four times using 1-mL pipette. Then, cell pellet was collected by centrifuging the tube at 200xg for 12 minutes at 25°C. To further stop papain reaction, cell pellet was resuspended with high-Ovomucoid solution (pH 7) containing 5 mg/mL of Trypsin inhibitor (Worthington Biochemical) and 5 mg/mL BSA (Sigma-Aldrich) in D-PBS. After centrifuging the tube at 200xg for 12 minutes at 25°C, cells were resuspended with panning buffer (0.02% of BSA and 5μg/mL of insulin in D-PBS). Mixed retinal cell suspension, which was passed through a sterile nylon mesh, was added to a negative panning dish coated with BSL-1 (5μg/mL, Vector Labs) for 30 minutes at room temperature. Then, cell suspension was transferred to another dish coated with BSL-1 (5μg/mL, Vector Labs) for 10 minutes. Further, cell suspension was transferred to positive panning dish coated with Thy1.2 (CD90, 1:800, Biorad) for 60-90 minutes at 37°C cell culture incubator. Before adding cell suspension to the positive panning dish, the dish was washed 9 times with D-PBS to remove azide. After Thy1.2 antibody panning, cell suspension was removed and the dish was washed 8 times with D-PBS. The dish was examined under microscope to make sure that only adherent cells remain. RGCs were detached from the dish by pipetting prewarmed RGC growth medium directly to the dish. Cells were collected by centrifuging the tube at 200xg for 12 minutes at 25°C. ~70,000 cells/well were seeded in a 8-chamber slide coated with either 10μg/mL rNPNT (R&D system), 2μg/mL BSA, or 1X Poly-D-Lysine (Sigma-Aldrich). All slides were treated with 1X Poly-D-Lysine for 1 hour at 37°C first and were stored at 4°C. Chamber-slide wells were coated with rNPNT or BSA for 2 hours at 37°C before seeding cells. Medium was changed every other day. For peptide treatment, medium was changed on the second day with RGC growth medium containing either 10μM GRGDSP or 10μM GRADSP. After 5 days, cells were fixed for immunocytochemistry.

RGC growth medium contains NeuroBasal A-Medium (Thermo Scientific), 5μg/mL Insulin (Sigma-Aldrich), 110μg/mL Sodium Pyruvate (Gibco), 100U/mL Penicillin-100μg/mL streptomycin (ThermoFisher), 1X SATO Supplement (100μg/mL BSA, 100μg/mL Transferrin (Sigma-Aldrich), 16μg/mL Putrescine (Sigma-Aldrich), 0.06μg/mL Progesteron (Sigma-Aldrich), and 0.04μg/mL Sodium selenite (Sigma-Aldrich)), 40ng/mL 3,3’,5-triiodo-L-thyronine sodium salt (T3, sigma-Aldrich), 292μg/mL L-glutamine (Thermos Scientific), 1X NS21 Supplement (R&D Systems), 1X B27 Plus (Thermos Scientific), 5μg/mL N-acetyl-L-cysteine (NAC, Sigma-Aldrich), 4.2μg/mL Forskolin (Sigma-Aldrich), 50ng/mL BDNF (Peprotech), and 10ng/mL ciliary NeeuroTrophic Factor (CNTF, Perotech).

### Electrophysiology

Visual response properties were determined as previously described (8), with minor modifications. Briefly, isoflurane-anesthetized adult mice were head-fixed and a 16-channel silicon multi-electrode (Neuronexus Technologies) was inserted into at an angle of 45 deg to the midline and 45 deg to the horizontal plane through a craniotomy located ~1.5 mm lateral to the midline and ~1.5 anterior to lambda. Electrodes were labeled with 1,1’-Dioctadecyl-3,3,3’,3’-Tetramethylindocarbocyanine Perchlorate (Invitrogen) and localization was confirmed *post hoc* via fluorescent microscopy. Multiunit signals were acquired at ~25 kHz and filtered between 0.7-7 kHz using a System3 Workstation (Tucker-Davis Technologies). Individual units were identified *post hoc* using independent components analysis. Visual stimuli were displayed on an LCD monitor subtending ~80 x 50 deg of visual space and placed directly in front of the animal. Stimuli consisted of drifting square waves presented at 12 different orientations and 6 different spatial frequencies, each presented 5-7 times. In addition, full-field flash and gray screens were presented to provide robust visual stimulus and determine spontaneous firing rate, respectively. Stimuli were presented in three ocular conditions: to both eyes together (both open, BO), with the ipsilateral eye covered (contralateral open, CO), and with the contralateral eye covered (ipsilateral open, IO). In each ocular condition, units were determined to be visually responsive if 1) the maximally-elicited mean firing rate was 3 standard deviations (SDs) greater than spontaneous, 2) the rate was 3 SDs greater than spontaneous in at least two-thirds of trials, and 3) the maximal mean rate was significantly different than the spontaneous rate, as determined by a Wilcoxon’s Rank Sum test. Based on their responsiveness under each condition, units were classified into one of seven potential types (9): monocularly-driven by the contralateral eye (BO^+^CO^+^IO^-^), monocularly-driven by the ipsilateral eye (BO^+^CO^-^IO^+^), binocular simple (BO^+^CO^+^IO^+^), binocular emergent (BO^+^CO^-^IO^-^), binocular inhibited by ipsilateral (BO^-^CO^+^IO^-^), binocular inhibited by contralateral (BO^-^CO^-^IO^+^), or binocular cross-inhibited (BO^-^CO^+^IO^+^).

### Looming assay

Looming stimuli were presented to the animals in white rectangular arena (47×37×30 cm) with an opaque shelter placed in a corner. Entrance of the shelter was facing the center of the arena. The arena was diffusely and evenly illuminated from above and was located within a light-proof and sound isolated room to maintain constant environmental conditions. Camera with frame rate of 30 FPS for capturing mouse’s behavior was secured to the stand next to the arena. For presenting the looming pattern, we put a computer monitor on top of the enclosure and second monitor with duplicated screen on table next to the arena, where the experimenter could control stimuli presenting and recording of animals. Looming object pattern was made in Photoshop as 25 shots of expanding black circle from 2 to 50° of visual field in 460 ms and remains 50° for an additional 540 ms. This 1sec sequence was repeated 10 times. Size of the black circle was calculated according the formula for visual angle calculation V=2 arctan (S/2D), where V is visual angle (i.e. 2-50°), S is size of the black circle and D is distance to the circle (i.e. 30 cm). Using this formula - “S” can be calculated as follows → S=2Dtan(V/2). All mice were tested only once to avoid a habituation to the looming stimulus. At the beginning of test, animals were let freely investigate the arena and the shelter for the period of 10 mins before the recording started. We started the video capturing approximately 10 sec prior to looming stimulus and the looming stimulus began when the animal was moving around the center of arena. Videos were recorded 10 sec prior, during and after looming stimulus. Then, animal’s behavior was scored manually during 10 sec of looming stimulus using ANY-maze software. Similar to (10), we scored 4 types of behavior - freezing, running, hiding and ambulation. Freezing was defined as period of one or more seconds in which the animal was completely immobile. Running was scored as activity in which the mouse started to move at least two times faster, then the average speed before stimulus onset. Hiding was defined as activity when the mouse was completely hidden in the shelter. Ambulation was defined as all other locomotor activity performed in the open arena.

### Prey capture

The task was recorded in a rectangular, white acrylic arena 47 cm long x 37 cm wide x 30 cm high using the ANY-maze software. The arena was diffusely and evenly illuminated from above and was located within a light-proof and sound isolated room to maintain constant environmental conditions. The floor of the arena was cleaned between each trial and mice with 30% EtOH and let to dry out completely. In order to perform this task, we followed the protocol from the study of (11). Similarly, each animal in our protocol underwent a 6-day acclimation protocol followed by testing day. The first three days of the protocol, mice were handled by the experimenter and exposed to crickets in their home cages overnight, along with their standard diet. On the third day, the mice were introduced to the testing arena environment for 10 min and their food deprivation started. Fourth day of the protocol was Day 1 of arena testing. Live crickets and mouse were placed together in the testing arena. Each mouse was given three 10-min trials to catch a cricket within the arena. If the mouse caught the cricket within these 10 min, the trial was scored as a capture success, and the capture time for that trial was recorded. The next trial was started after cleaning with 30% EtOH and drying out the arena. If a cricket was not captured within 10 min, the mouse was removed from the arena for 1 min and returned into the arena with a new cricket to start the next trial. At the same time, the mean capture time and average speed during the hunt for each mouse on each day was calculated. After the three trials, each mouse was offered free access to mouse chow and two live crickets in their home-cage for approximately 6 hours before starting next food deprivation. Training of prey-capture and food deprivation was continued for three days. By day 3 of hunting in the arena, 98% of mutant mice and 100% of control mice achieved 100% capture success. The 6 days of acclimation was followed by the Testing day when mean capture time and average speed during the hunt was measured. All tested animals achieved 100% capture success on testing day.

---

**Figure S1.**
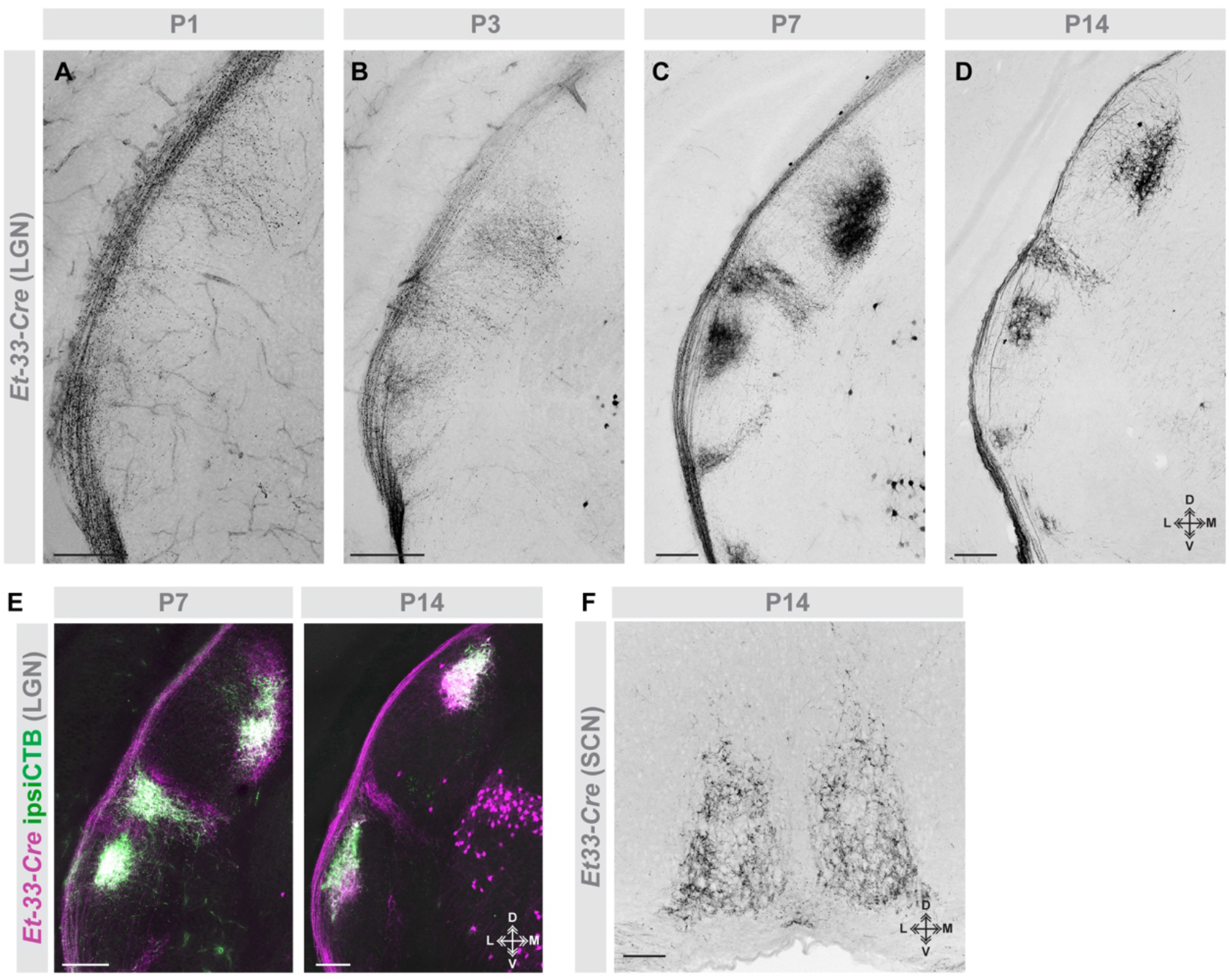
*Et33-Cre* labels central projections of ipsiRGCs during postnatal development. **A-D.** Genetically labeled ipsiRGCs innervate the developing lateral geniculate nucleus (LGN) in *Et33-Cre::Rosa-Stop-tdT* mice. Arrows highlight that *Et33-Cre* labels a small population of cells in dorsal thalamus. **E.** Monocular injection of anterograde tracer cholera toxin B (CTB) co-labels genetically labeled ipsiRGCs in *Et33-Cre::Rosa-Stop-tdT* mice. **F.** Genetically labeled ipsiRGCs innervate the suprachiasmatic nucleus (SCN) in *Et33-Cre::Rosa-Stop-tdT*mice. Scale bars in A,B = 200 μm; in B = 50 μm; in C = 50 μm for C,D; in E = 200 μm; in F = 100 μm.

**Figure S2.**
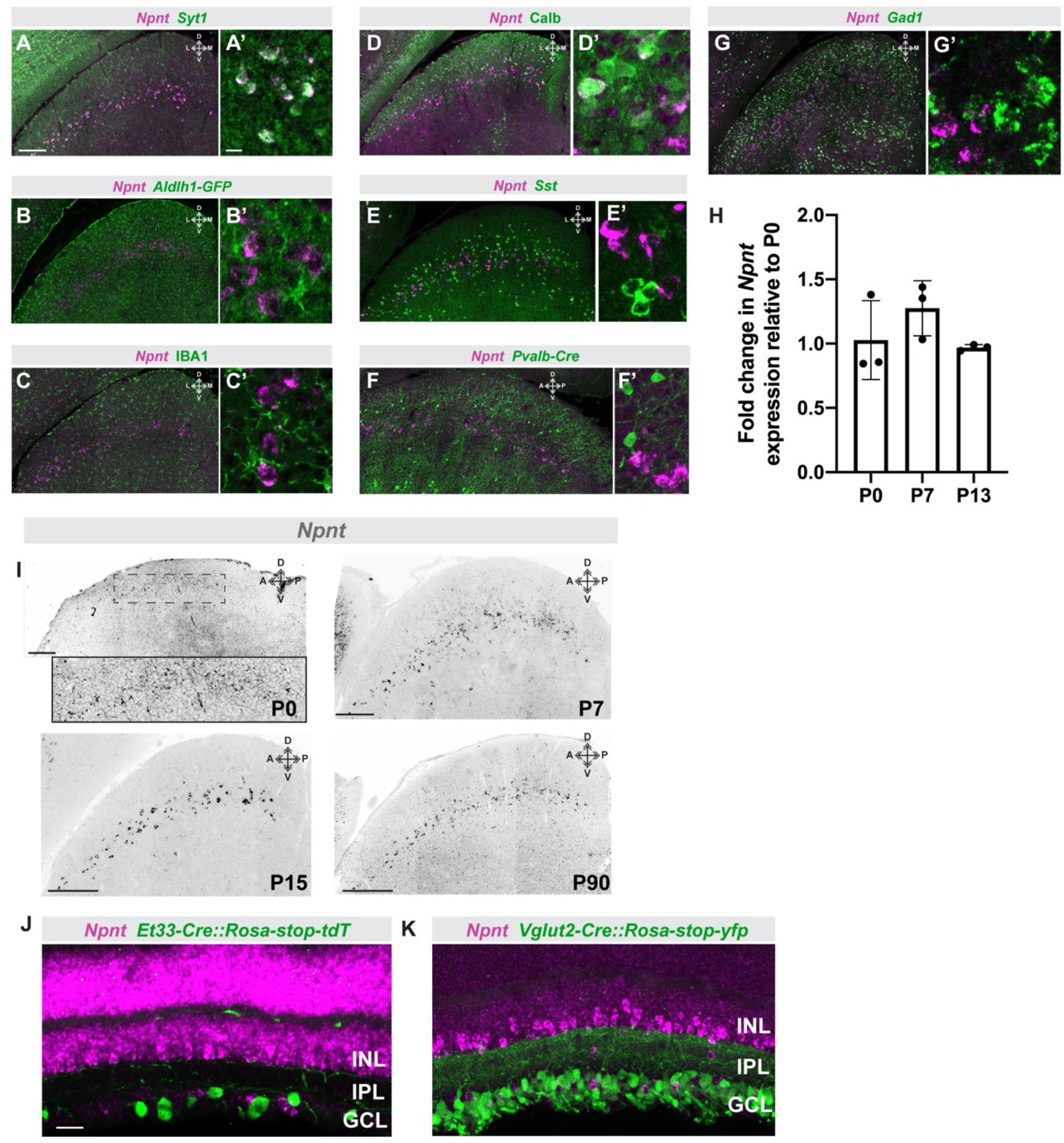
*Npnt* is expressed by glutamatergic neurons in SC and not by RGCs in retina. **A-G**. Co-labeling *Npnt^+^* cells (by *in situ* hybridization) with multiple SC cell type markers (genetically, by *in situ* hybridization, or by immunolabeling), specifically with glial markers *Aldh1l1* and IBA1 and neuronal markers *Syt1*, Calb, *Sst, Pvalb*, and *Gad1* at P15. **H**. *Npnt* mRNA expression in the postnatal SC detected by qPCR. Bars represent means +/− SD (N=3 mice). **I**. Labeling of *Npnt^+^* cells (by *in situ* hybridization) in the P0, P7, P15, and P90 SC. Inset shows high magnification in *Et33-Cre::Rosa-Stop-tdT*. **J-K**. Labeling *Npn^+^* cells in retina (by *in situ* hybridization) and co-labeling of ipsiRGCs in *Et33-Cre::Rosa-Stop-tdT* (J) or all RGCs in *Vglut2-Cre::Thy1-Stop-yfp* (K) reveals that *Npnt^+^* cells are not expressed by RGCs. GCL - ganglion cell layer; IPL - inner plexiform layer; INL - inner nuclear layer. Scale bar in A = 200 μm and inset = 20 μm for A-G; in I =200 μm; in J,K = 40 μm.

**Figure S3.**
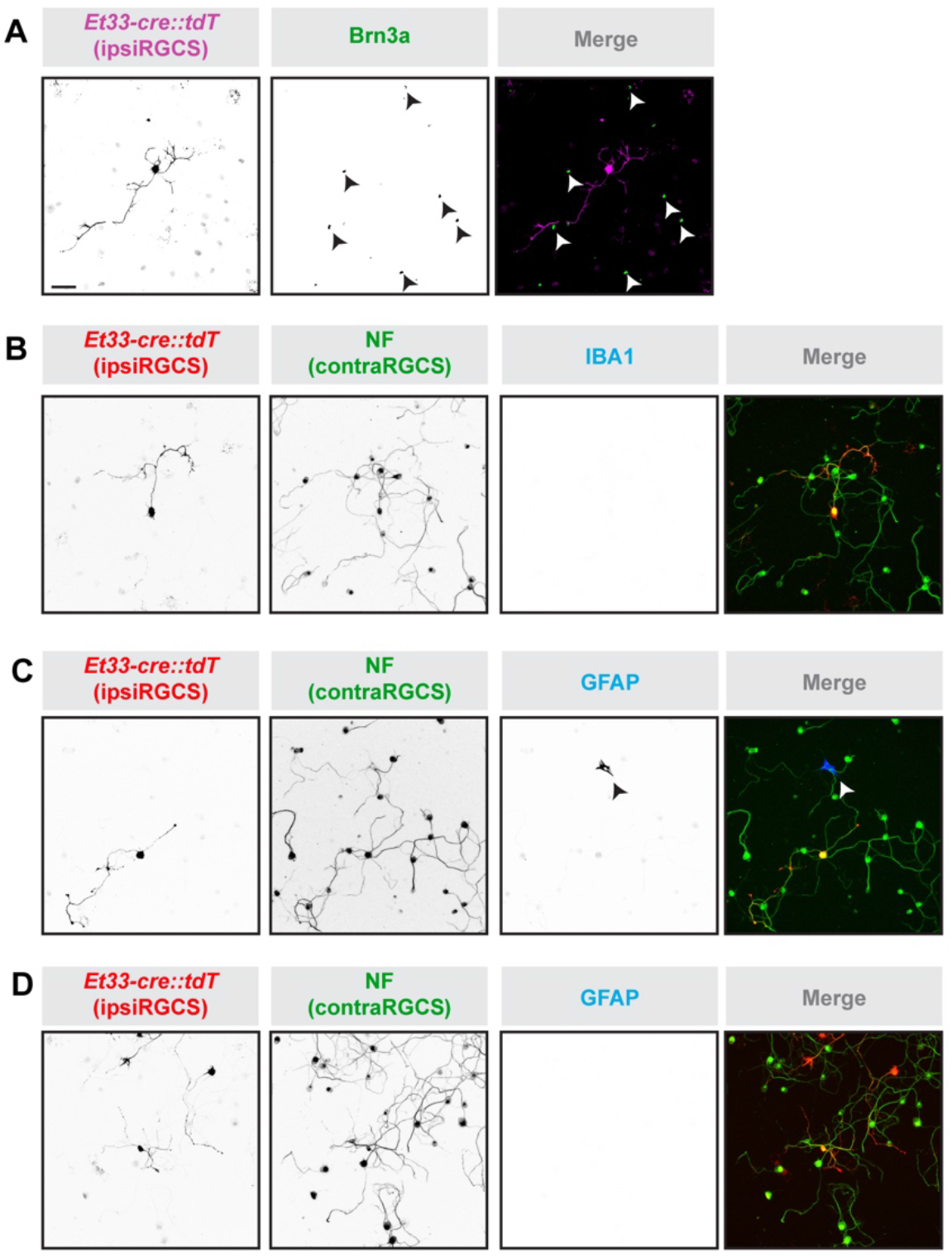
Purity of immunopanned RGC cultures. **A-D**. Genetically labeled ipsiRGCs cultured from *Et33-Cre::Rosa-Stop-tdT* mice do not co-express Brn3a (**A**), IBA1 (**B**), or GFAP (**C,D**). Cells immunoreactive to glial markers were rare to find in these cultures. One rare occurrence is shown in (**C**). Scale bar in A = 100 μm for A-D.

**Figure S4.**
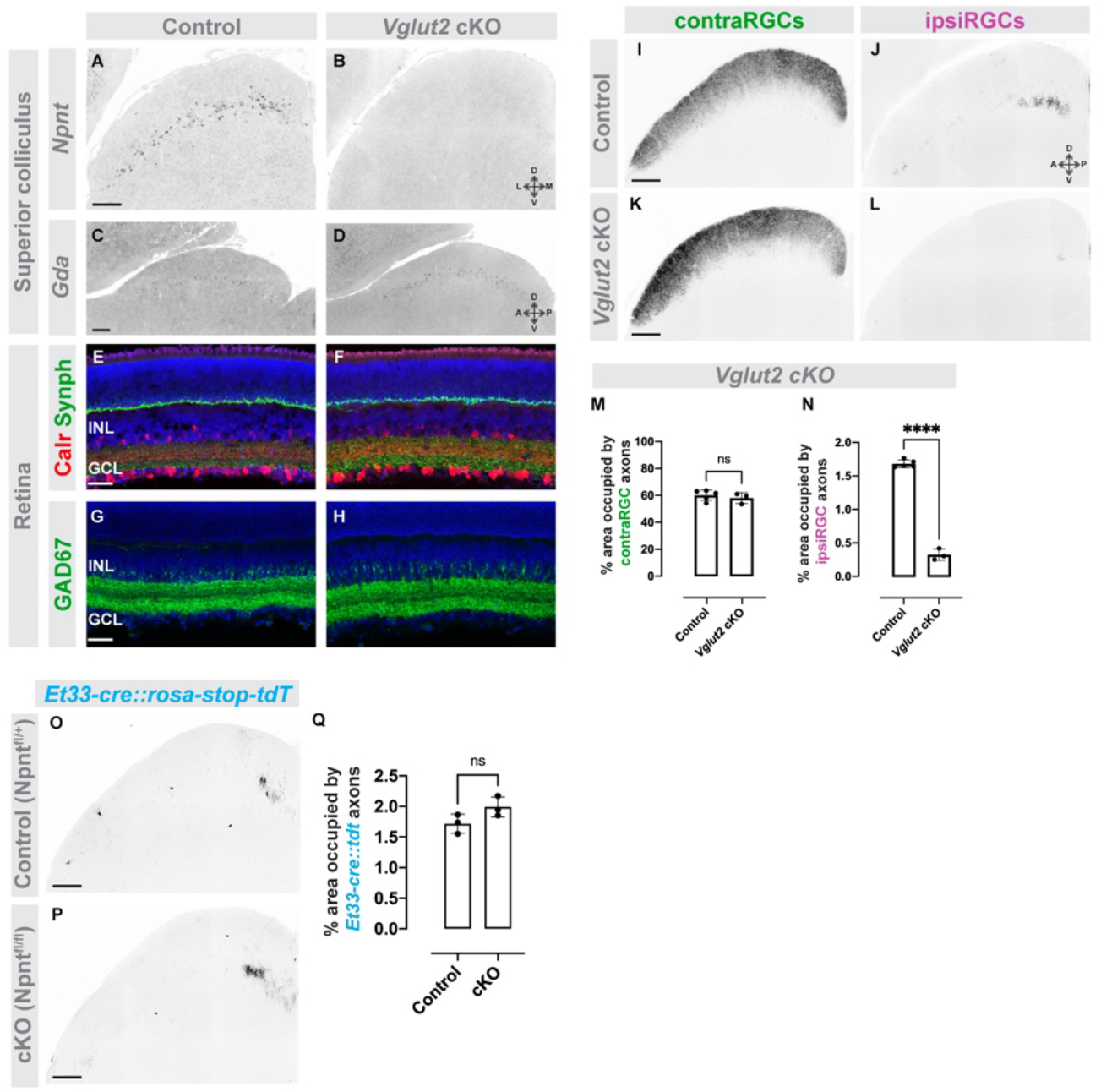
Conditional deletion of *Npnt* from *Vglut2^+^* neurons does not affect retinal or collicular cytoarchitecture and leads to a loss of ipsiRGC inputs to SC. **A,B.** Absence of *Npnt* mRNA expression (detected by ISH) after conditional deletion of deletion of *Npnt* from *Vglut2^+^* neurons in *Vglut2-Cre::Npnt^fl/fl^* mice (B; *Vglut2* cKO). **C,D**. ISH of *Gda* mRNA shows that SC cells in the deepest lamina remain unaffected after conditional deletion of deletion of *Npnt* from *Vglut2^+^* neurons in *Vglut2* cKO (D) mice. **E-H**. Cytoarchitecture of retina remains unaffected after conditional deletion of deletion of *Npnt* from *Vglut2^+^* neurons in *Vglut2* cKO (F) mice. **I-L**. CTB-labeling of contra- and ipsiRGCs projections to SC show that ipsiRGC axons are absent in *Vglut2* cKO mice. **M,N**. Quantification of the area of SC occupied by contra- (G) and ipsiRGC (H) projections in E,F. Bars represent means +/− SD. **** indicates P<0.0005 by Student’s t-test (N=3 mice). **O,P**. Projections of genetically labeled ipsiRGCs to SC are unaffected by the conditional deletion of *Npnt* from *Et33^+^* ipsiRGCs in *Et33-Cre:Npnt^fl/fl^::Rosa-Stop-tdT* mice. **Q**. Quantification of the area of SC occupied by genetically labeled ipsiRGC projections in I,J. Bars represent means +/− SD. No significant difference detected by Student’s t-test (N=3 mice). Scale bar in A,B,E,F,I,L,M = 200 μm; in C,D = 100 μm.

**Figure S5.**
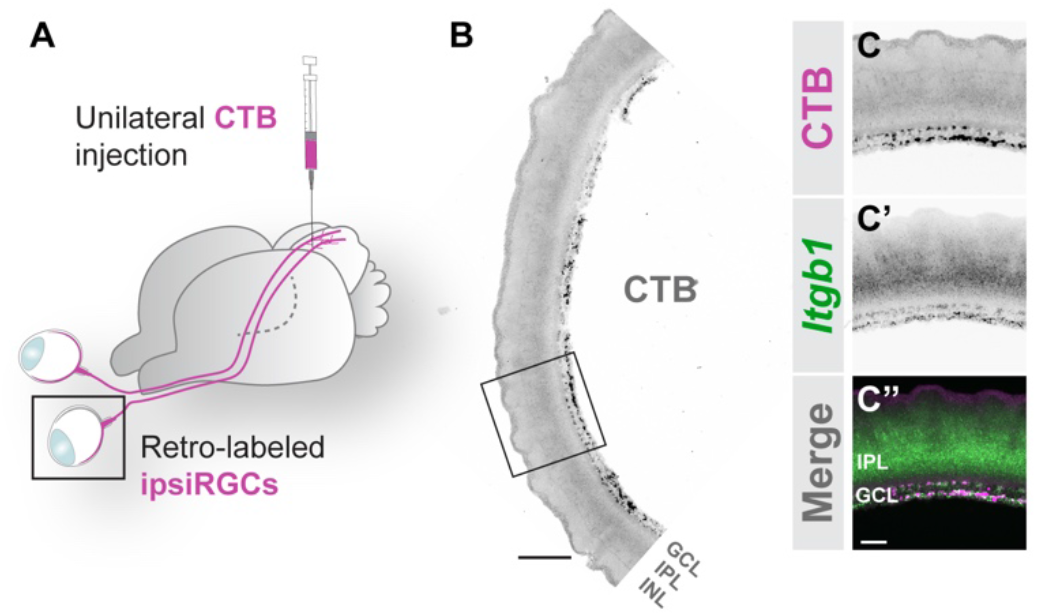
Retro-labeled SC-projecting ipsiRGCs express *Itgb1* mRNA. **A**. Schematic of retrograde labeling of ipsiRGCs with intracollicular injection of CTB. **B, C**. ISH for *Itgb1* mRNA in P14 retinal cross sections following retrograde labeling of ipsiRGCs with CTB. **B** depicts a low magnification image representing the ventrotemproal crescent of retina; **C** depicts a high magnification image of *Itgb1* mRNA and CTB labeled ipsiRGCs. Scale bar in B = 150 μm; in C = 40 μm.

**Figure S6.**
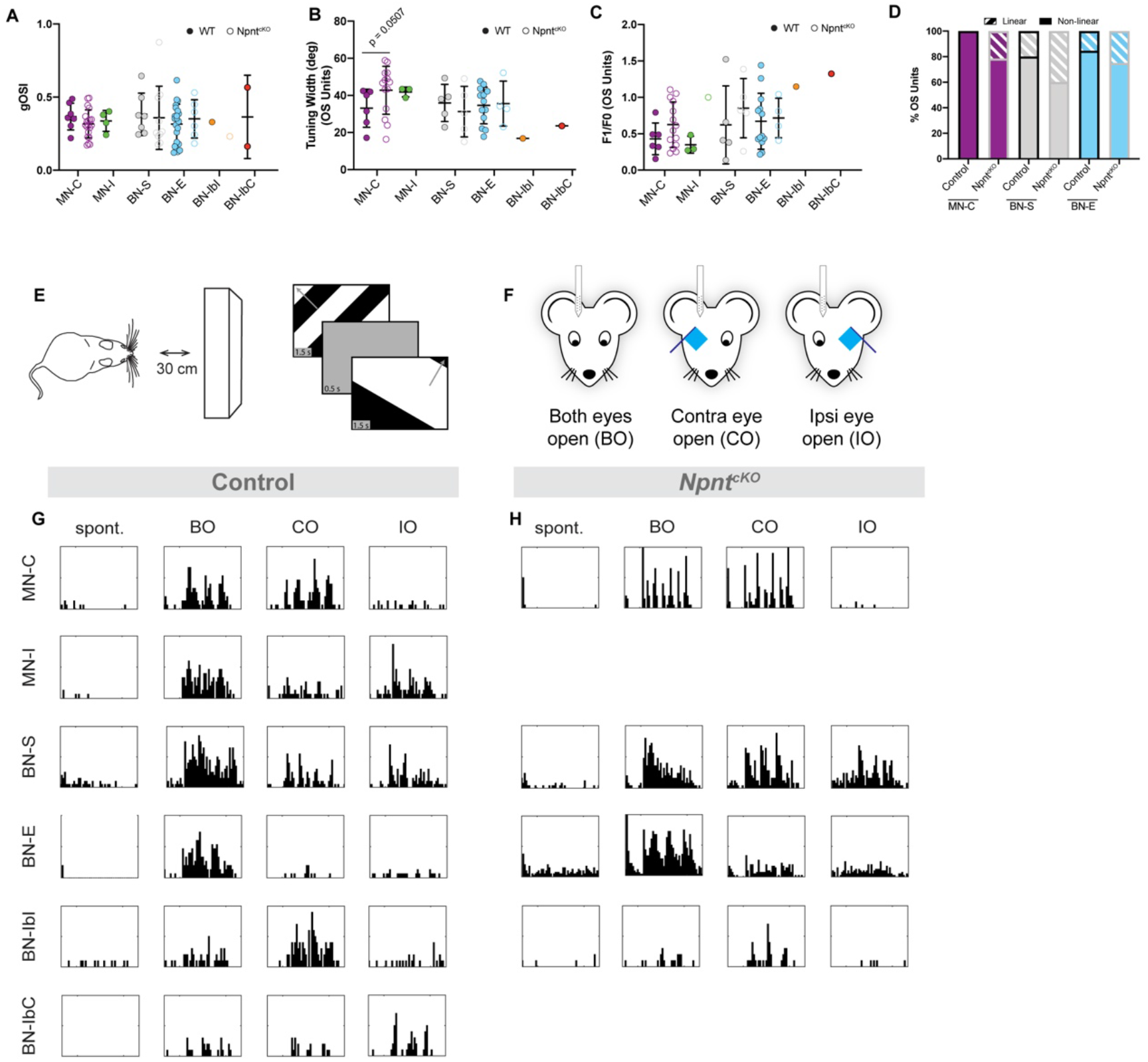
Classification of monocular and binocular subtypes in the SC. **A-D.** Quantification of the global orientation-selectivity index (gOSI) (**A**), tuning width of orientation-selective (OS) units (**B**), and F1/F0 ratio of OS units (**C**) in the indicated subtypes of visual neurons in control (closed circles) and *Npnt-cKO* (open circles) mice. (**D**) Proportions of OS units exhibited linear (checked) and non-linear (solid) summation of spatial stimuli based on F1/F0 ratio in the indicated subtypes of visual neurons in control (black border) and *Npnt-cKO* (gray border) mice. MN-C: monocularly-modulated contra-driven; MN-I: monocularly-modulated ipsi-driven; BN-S: binocularly-modulated simple; BN-E: binocularly-modulated emergent; BN-IbI: binocularly-modulated inhibited by ipsi; BN-IbC: binocularly-modulated inhibited by contra. **E,F.** Schematic of visual stimulus paradigm in which drifting gratings were presented directly in front of mice (**E**) under three ocularity conditions (**F**). **G,H.** Representative peri-stimulus spike time histograms from indicated subtypes of neurons identified in control (**G**) and *Npnt-cKO* (**H**) mice.

